# Loss of the tumor suppressor PTEN activates cell-intrinsic interferon signaling to drive immune resistance

**DOI:** 10.1101/2025.09.17.676669

**Authors:** Yubao Wang, Cherubin Manokaran, Jiajia Chen, Hao Gu, David Liu, Philippe Bousso, Jean J Zhao, Thomas M Roberts

## Abstract

Loss of the tumor suppressor PTEN is strongly associated with a lack of response to immune checkpoint blockade therapies in cancer patients, but the underlying mechanisms are not fully understood. We have developed a transformation model where knocking out PTEN in human mammary epithelial cells drives dependence on the p110β subunit of PI3K and, notably, elicits a robust induction of endogenous retroviral elements (ERVs) and activation of the interferon signaling. This constitutive cell-intrinsic interferon response, hallmarked by hyperactivated STAT1, is also observed in human tumors with a PTEN-low status. We further find that PTEN deficiency renders cancer cells resistant to the cytotoxic effects of immune cells and interferon-γ. Notably, PTEN loss also results in a dependency on an activated DNA damage response pathway, leading to an exquisite vulnerability to CDK12 inhibition. Our study suggests an interferon adaptation model in which tumors driven by PTEN deficiency inherently activate the interferon response, enabling them to adapt to interferon cytotoxicity and gain resistance to immunotherapies.

## Introduction

The tumor suppressor gene *PTEN* was initially identified due to its widespread loss-of-function alterations in human cancer^1–3^, and was then found to encode a protein with lipid phosphatase activity that antagonizes the action of phosphatidylinositol 3-kinases (PI3Ks)^4,5^. Both *PTEN* and *PIK3CA*, which encodes p110α□□one of the catalytic subunits of Class I PI3K, are among the approximately one dozen genes most frequently mutated in human cancer^6^. Interestingly, genetic alterations in *PTEN* tend to be mutually exclusive with those in *PIK3CA*/p110α^7^ (Fig. S1A). Studies have found that PTEN-deficient human cancer cell lines often rely on another catalytic subunit of PI3K, PIK3CB/p110β, to maintain elevated PI3K activity and tumorigenic potential^8–10^. However, recent studies have demonstrated that PTEN-deficient tumors do not invariably rely on PIK3CB^11–14^, posing a challenge to developing PIK3CB-targeted cancer therapies.

Alongside studies focused on developing targeted therapies, a new paradigm in cancer treatment involves stimulating anti-tumor immunity. A significant obstacle in immunotherapy is that only a subset of cancer patients responds to the treatment, and the mechanisms of resistance in most resistant patients remain unclear. Notably, *PTEN* mutation has been found robustly associated with tumor resistance to immune checkpoint blockade in various cancer types, including melanoma^15^, metastatic uterine leiomyosarcoma^16^, glioblastoma^17^, metastatic triple-negative breast cancer^18^, and microsatellite instability-high/mismatch repair-deficient gastrointestinal tumors^19^. For instance, melanoma patients with PTEN loss are approximately six times more likely to be resistant to checkpoint blockade with antibodies directed against cytotoxic T lymphocyte antigen-4 (CTLA-4) and programmed death receptor-1 (PD-1)^20.^

These observations and advances raise several key questions regarding malignancies driven by PTEN inactivation. First, do cancers lacking or with low levels of PTEN expression significantly rely on PIK3CB/p110β for growth? Secondly, how does PTEN loss lead to a lack of response to immune checkpoint blockade (ICB) therapies? In particular, is the insensitivity to immunotherapies due solely to a potential immune suppressive microenvironment that was documented in PTEN-deficient tumors^15,21,22^? Alternatively, are PTEN-deficient tumor cells intrinsically resistant to cytotoxic immune cells? In this study, we developed a unique transformation model to demonstrate that PTEN loss drives both PIK3CB/p110β dependence and cell-intrinsic activation of the interferon signaling. Our study suggests that tumor-intrinsic interferon signaling, through an adaptative mechanism to interferon cytotoxicity, is a potential mechanism of resistance to immunotherapies.

## Results

### PTEN-low human cancer cells depend on PIK3CB, the p110β catalytic subunit of PI3K, for growth and survival

Although studies in genetically engineered mouse models and human cancer cell lines have highlighted the importance of PIK3CB/p110β for the growth of PTEN-deficient tumors, the results have been controversial^8–14,23^. To assess the robustness of PIK3CB dependence in cancers with loss of or low expression of PTEN, we analyzed recently generated omics datasets to study the association between PTEN status and PIK3CB dependence in a large number of human cancer cell lines. We first found that *PTEN* mRNA levels or genetic alterations are not the most robust marker for determining PTEN protein status (Fig. S1B). Instead, the protein abundance of PTEN itself faithfully reflects PI3K activity (Fig. S1C-F).

Using PTEN protein abundance as a molecular marker, we found that, among 320 anticancer drugs tested in 482 cancer cell lines in GDSC database^24^, the top positive correlates are AZD6482 (also named KIN193), a PI3K p110β catalytic subunit-specific inhibitor, and TGX-221, the chemical predecessor to AZD6482/KIN193 (Fig. 1A). In other words, the less PTEN protein cancer cells express, the more sensitive they are to p110β inhibition. Further strengthening the association, across proteins in the RPPA dataset PTEN abundance was most correlated to cancer cell sensitivity to AZD6482 (Fig. S2A). Notably, the strong association with p110β inhibitors could have been under-estimated if other molecular markers of PTEN status, such as PTEN copy number, mutation status, or mRNA expression, were used for the analysis (Fig. S2B).

**Figure 1.**
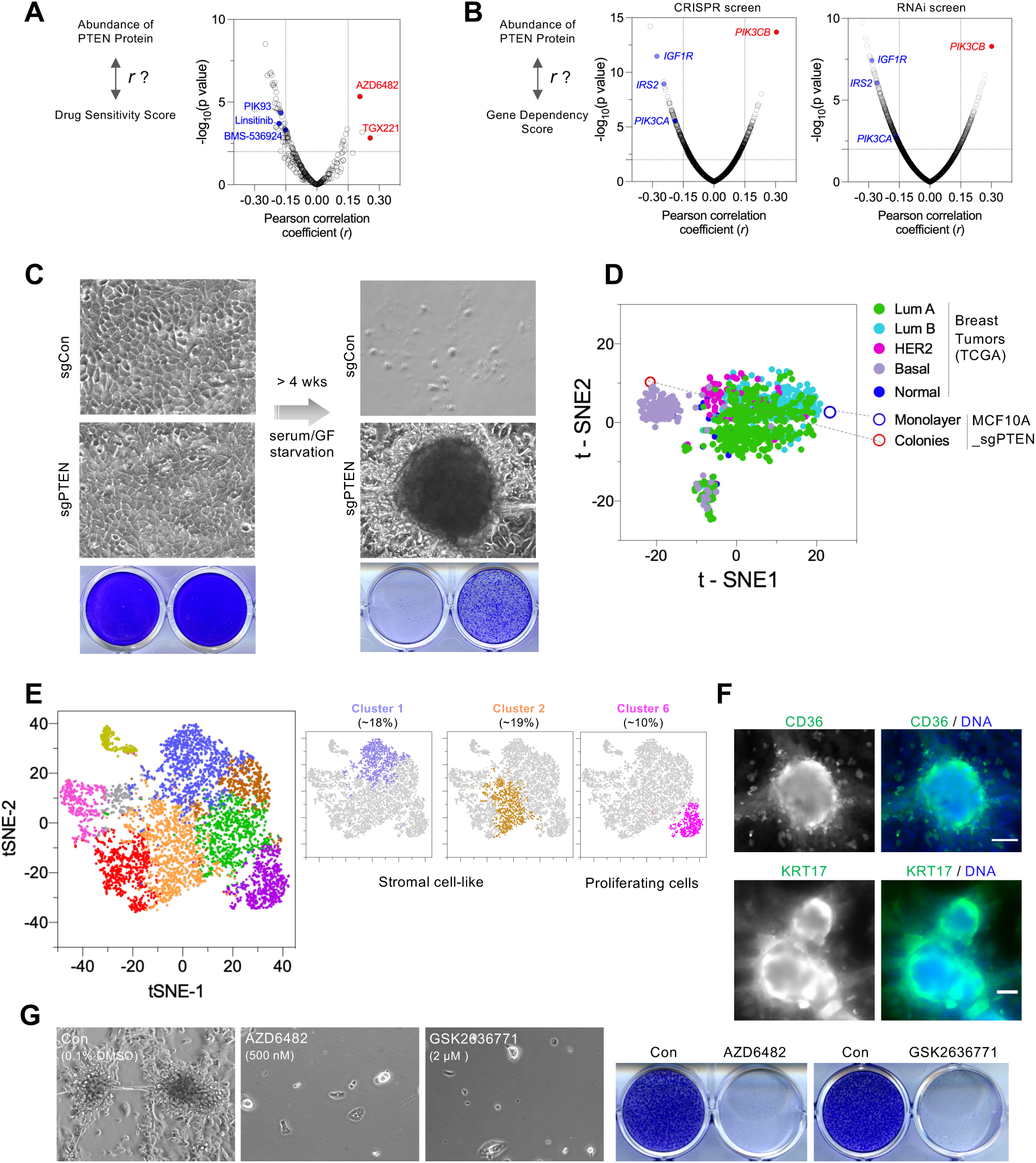
A PTEN loss-driven cellular transformation model. (A) Correlation of PTEN protein abundance with anticancer drug sensitivity, illustrated by the diagram (Left). The analysis (Right) was based on GDSC and RPPA datasets; in total, 482 cancer cell lines have PTEN protein abundance data as well as drug response data (n = 320, including clinical, pre-clinical, and tool compounds). Pearson correlation coefficient (x-axis) for each individual compound and the statistical significance of the correlation (y-axis) are plotted. (B) Pearson correlation analysis between PTEN protein abundance and gene dependency, illustrated by the diagram (Left). The analysis was performed using the genome-wide Avana library CRISPR screen (Middle) and Project DRIVE RNAi screen (Right). Pearson correlation coefficient (x-axis) for each individual gene and the statistical significance of the correlation (y-axis) are plotted. (C) An *in vitro* model of cellular transformation. MCF10A cells stably transduced with control (sgCon) or PTEN-targeting guide (sgPTEN) were grown to confluence, imaged, and fixed (Left), while replicate cells were cultured in the absence of serum or growth factors for one month (Right). Bright-field images and crystal violet staining are shown. (D) A gene signature was derived for the sgPTEN colonies and used to generate a t-SNE plot of all samples in the TCGA breast cancer cohort. sgPTEN colony and monolayer samples were included in the analyses to determine how they cluster with the patient data. PAM50 subtypes are indicated. (E) Transformed sgPTEN cells were subjected to single-cell RNA-seq, and a t-SNE projection of cells colored by automatic clustering is shown. Clusters of cells with stromal and proliferative markers are highlighted. (F) Immunostaining of transformed sgPTEN cells with anti-CD36 (Top) and anti-KRT17 (Bottom). Note that colonies were surrounded by CD36- and KRT17-positive cells (G) MCF10A_sgPTEN cells were subjected to deprivation of serum and growth factors in the presence of vehicle or the indicated PIK3CB inhibitors. After one month of culture, cells were imaged (Left), fixed, and stained with crystal violet for whole-well views (Right).

We then analyzed CRISPR screen data to identify which gene(s) may be required for the growth of PTEN-low cancer cells. In a genome-wide screen using 789 cancer cell lines^25^, PIK3CB/p110β was the leading positive correlate with PTEN protein abundance among 18,119 genes tested in the screen (Fig. 1B, S2C). Thus, the less PTEN protein cancer cells express, the higher dependence they have on PIK3CB/p110β□for cell proliferation. In contrast, PIK3CA/p110α□dependence is a significant, negative correlate of the PTEN protein signal (Fig. 1B, S2C). We further analyzed other loss-of-function screens^26,27^, and observed a strong positive association between PTEN protein abundance and PIK3CB/p110β gene dependence (Fig. 1B, S2D). Therefore, PIK3CB/p110β appears to be an exceptionally strong dependency in cancer cells with a PTEN-low status.

Just as PIK3CB appears to be a compelling dependency in this context, we sought to determine whether PTEN protein is the chief determinant of PIK3CB/p110β dependency.

Among 214 proteins tested in the RPPA assay, PTEN protein abundance showed the strongest association (Fig. S2E-F). Therefore, low PTEN protein is not only associated with PIK3CB/p110β dependency but it is also likely the strongest determinant or predictor of this dependence.

### Ablating PTEN fails to confer an immediate dependence on PIK3CB/p110β in untransformed cells

We then asked if restoring PTEN expression in PTEN-deficient cancer cells would alter their dependence on p110β. Cancer cells stably transduced with PTEN were able to grow following an extended period of culture while displaying impaired growth in comparison to control cells (Fig. S3A). PTEN-expressing cells had substantially reduced phosphorylation of AKT and its substrate PRAS40 (Fig. S3B), despite increased expression of insulin receptor substrates (IRSs) especially IRS2^28^. While control cells had an expected dose response to AZD6482/KIN193 in terms of AKT phosphorylation (*i.e.*, 1 □M dose is capable of reducing AKT phosphorylation by >80%), the residual phospho-AKT signal in cells with ectopic PTEN expression was more resistant to p110β inhibition (Fig. S3C, S3D). In contrast, the phospho-AKT signal in PTEN addback cells became more sensitive to p110α inhibition (Fig. S3C, S3E). These data suggest that the loss of PTEN protein is essential for p110β dependence of PI3K signaling.

We next used CRISPR/Cas9-mediated gene editing to ask if reducing PTEN expression in PTEN-proficient untransformed cells (human mammary epithelial cells, HMECs) would engender PIK3CB dependence. Consistent with an efficient loss of PTEN protein in cells transduced with PTEN-targeting guide (sgPTEN), these cells demonstrated elevated PI3K/AKT signaling, indicated by AKT, PRAS40, and S6 phosphorylation, as well as the electrophoretic mobility shift of 4E-BP1. However, despite the almost, complete loss of PTEN protein, the cells were not more sensitive to p110β inhibition (Fig. S4A). Thus, we concluded that the knocking down PTEN expression alone may not be sufficient for cells to gain the p110β dependence that is often observed in tumor-derived cell lines with naturally occurring PTEN-low status.

### PTEN loss-induced transformation drives PIK3CB/p110β dependence

Seemingly, there is a discordance between the effect of knocking down PTEN expression in untransformed cells (*i.e*., no immediate gain of sensitivity to PI3K/p110β inhibition) and the top p110β□dependency observed in PTEN-low cancer cells. We hypothesized that PTEN-low status is one integral aspect of transformation in the course of tumor formation, and that cells subsequently adapt to the loss of PTEN and become dependent on PI3K/p110β. Guided by this hypothesis, we asked if we could establish a PTEN loss-induced transformation model *in vitro*, and then use the model to study if and how p110β dependence arises.

Transformation *in vitro* has been extensively modeled by using rodent fibroblasts in foci assays, in which contact inhibition of cell growth is lost upon the introduction of a single potent oncogene or the loss of a tumor suppressor. In human epithelial cells, acquisition of the ability to grow in anchorage independent conditions is often used as a surrogate for transformation, with a combination of multiple genetic lesions required for the cell transformation^29^. We chose MCF-10A, a spontaneously immortalized human epithelial cell line that is widely used for transformation assays^30^. We reasoned that the loss of PTEN or activation of PI3K/AKT could be an early event in tumor formation that enhances nutrient uptake and cell growth under growth factor-limiting conditions resembling environment for tumor genesis *in vivo*.

Thus, we tested various starvation conditions and found that loss of PTEN is able to support cell survival and morphological transformation following long-term starvation, a condition under which all control cells perished (Fig. 1C). Along with striking changes of cell morphology (*i.e*., from a uniform monolayer to a morphologically heterogeneous cell population consisting of visible colonies), gene expression profiling of sgPTEN monolayer cells and sgPTEN colonies revealed a profound difference of gene expression between the two states. Notably, comparing their gene expression with that of human breast tumors found that while the sgPTEN monolayer shared its gene expression pattern with luminal A breast cancer, sgPTEN colonies were more similar to basal-like breast cancer (a molecular subtype known to have the most frequent PTEN loss among all breast cancer subtypes^7^) (Fig. 1D). We further performed single-cell mRNA sequencing (scRNA-seq) and found that cells from the transformed colonies clustered into six distinct populations with gene expression characteristic of different cells types including stromal-like cells, myoepithelial cells, perivascular-like cells, and proliferative cells (Fig. 1E, S4B). In line with the heterogeneity in gene expression, the transformed cells displayed morphologically distinct cells, with colonies surrounded by cells positive for the expression of CD36 and Keratin 17 (Fig. 1F, S4C). Notably, CD36 was recently identified as a marker for progenitor cells in mammary basal epithelium^31^, while Keratin 17 has been known to be expressed in a specific sub-cluster of human mammary basal epithelial cells^32^. These observations suggest that this transformation model recapitulates some key characteristics of human breast cancer, with an overall gene expression landscape close to basal-like breast cancer and marked by a heterogeneous group of different cell types and states.

Using this model, we then studied how the resulting cells respond to PI3K inhibition. Interestingly, while PI3K signaling in both control cells and sgPTEN monolayers prior to transformation was apparently insensitive to PIK3CB/p110β inhibition, sgPTEN cells in the resulting colonies were extremely responsive, *i.e*., phosphorylation of AKT and PRAS40 in colonies was completely abolished by AZD6482/KIN193 treatment (Fig. S4D). Furthermore, cell growth and survival under these conditions was also potently suppressed by p110β inhibition (Fig. 1G). These results thus suggest that PIK3CB/p110β dependence is acquired when epithelial cells are transformed by PTEN loss.

### Activation of interferon signaling during PTEN loss-induced transformation

We further studied the gene expression in PTEN loss-driven transformed cells compared with their pre-transformed counterparts. To our surprise, the transformed cells demonstrated a robust enrichment of interferon (IFN) signaling gene sets (Fig. 2A, S5A-B). Consistent with these observations in RNA-seq, quantitative RT-PCR analysis revealed an approximately ten-fold or higher induction of interferon-stimulated genes (ISGs), including those encoding a core transcription factor of the IFN pathway (STAT1), a nucleic acid sensor RIG-I/DDX-58, effectors and/or regulators of IFN signaling (USP18, ISG15, OAS3, OASL), as well as proteins essential for antigen processing and presentation (HLA-A, HLA-B) (Fig. 2B). Notably, the proteins of STAT1, STAT2, USP18, MDA5, and RIG-I became readily observed in PTEN loss-transformed cells, but nearly absent in cells prior to transformation (Fig. 2C).

**Figure 2.**
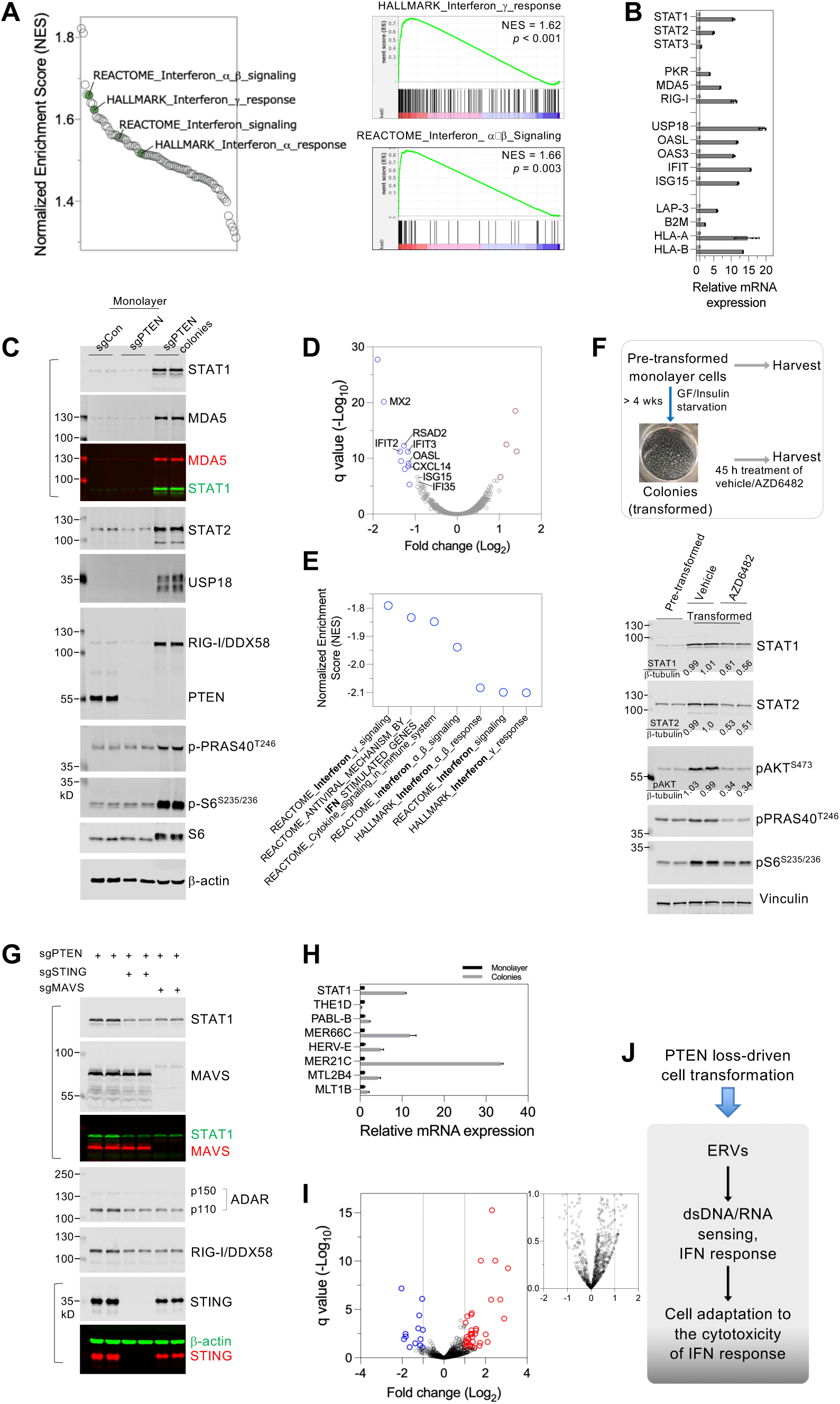
Activation of the interferon response PTEN loss-driven transformation. (A) (Left) Summary plot of normalized enrichment scores (NES) for gene sets significantly upregulated in transformed MCF10A_sgPTEN colonies compared to their monolayer counterparts in the pre-transformed state. Selected gene sets related to interferon signaling are noted. (Right) GSEA plots of indicated interferon signatures showing gene expression changes. NES and p-values are shown. (B) RT-qPCR analysis of indicated interferon-stimulated genes (ISGs) in transformed MCF10A_sgPTEN colonies (dark gray bars) versus their monolayer pre-transformed counterparts (light gray bars). Data are shown as mean ± SD (n = 3). (C) Fluorescent immunoblotting of the indicated whole-cell lysates. Merged blots are shown for selected membranes. The molecular weight of fluorescent protein markers is indicated. (D) Volcano plot showing differential gene expression in transformed MCF10A_sgPTEN cells treated with PIK3CB inhibitor GSK2636771 (2 μM) compared to vehicle control. MCF10A_sgPTEN transformed colonies were treated with PIK3CB inhibitor GSK2636771 (2 μM) or vehicle (0.1% DMSO, v/v) for 22 hours, followed by total RNA extraction and sequencing. Genes with absolute expression changes greater than 2-fold and q-value < 0.1 are colored blue (downregulated) or red (upregulated). Selected interferon-stimulated genes (ISGs) are noted. (E) Summary plot showing normalized enrichment scores (NES) for gene sets significantly downregulated in cells treated as in (D) (p-value < 0.05). (F) Fluorescent immunoblotting of whole-cell lysate from MCF10A_sgPTEN monolayer cells or derived colonies treated with vehicle or AZD6482/KIN193 for 45 hours. Quantification of indicated blots was performed using Image Studio (LI-COR Biosciences), with the signal normalized to vehicle-treated samples. (G) Fluorescent immunoblotting of whole-cell lysates from colonies derived from MCF10A_sgPTEN parental cells, MCF10A_sgPTEN/sgSTING clonal cells, or MCF10A_sgPTEN/sgMAVS clonal cells. Merged blots are shown for selected membranes. (H) RT-qPCR analysis of *STAT1* and the indicated ERVs in transformed MCF10A_sgPTEN cells, normalized to their pre-transformed monolayer counterparts. Data are shown as mean ± SD (n = 3). (I) Volcano plot showing differential expression of ERVs in MCF10A_sgPTEN colonies compared with monolayer cells. Significantly altered transcripts (absolute fold change > 1, q < 0.1) are colored red (upregulated) or blue (downregulated). A zoomed-in plot (top right) shows ERVs transcripts with less significant expression change (absolute fold change < 1 or q > 0.1). (J) A diagram illustrates that ERVs transcripts trigger dsDNA/RNA sensing and induce the interferon response during PTEN loss-driven transformation. As a consequence, transformed cells adapt to the cytotoxic effects of this chronic interferon response and are likely resistant to IFNs secreted by cytotoxic immune cells.

To understand how IFN signaling is activated during the transformation process, we first asked whether PIK3CB/p110β may be involved. To this end, we treated the transformed cells with a PIK3CB/p110β inhibitor for 16 hours (a period that was brief enough to avoid any discernible effects on cell viability) and then extracted total RNA for sequencing. Compared with vehicle treatment, p110β inhibition does not elicit an extensive alteration of gene expression, with only a few dozen genes showing expression changes greater than one-fold (Fig. 2D). This data is not entirely surprising, because p110β is not directly implicated in gene expression control but, rather, lies upstream of the signaling pathways regulating cell growth and survival. However, genes that are most susceptible to the treatment are heavily represented by ISGs (Fig. 2D), and genes downregulated by PIK3CB/p110β inhibition display a top-ranked enrichment in IFN gene sets (Fig. 2E). Consistently, PIK3CB/p110β inhibition in the transformed cells led to approximately half-reduction of both STAT1 and STAT2 proteins (Fig. 2F).

We hypothesized that PTEN loss in the context of chronic growth factor starvation may induce an abundance of endogenous double-stranded (ds) nucleic acids, which are subsequently sensed by the cells to initiate an interferon response. To test this hypothesis, we used CRISPR/Cas-mediated gene editing to obtain knockout clones of cells lacking STING (stimulator of interferon genes, a key protein in the dsDNA sensing pathway), or MAVS (mitochondrial antiviral-signaling protein, a dsRNA sensor). Both sgPTEN;sgSTING and sgPTEN;sgMAVS cells were able to form colonies under long-term growth factor starvation, but in each case colonies displayed reduced expression of ISGs, including STAT1, ADAR, and RIG-I/DDX58 (Fig. 2G). In line with a role of MAVS in mediating the IFN response in sgPTEN transformed cells, we also found that the expression of endogenous retrovirus (ERV) genes, including those known to trigger MAVS-mediated IFN response^33^, was induced in the transformed cells (Fig. 2H, 2I). These data suggest a model in which endogenous retroviral transcripts are involved in the activation of dsDNA/RNA-mediated IFN signaling during the process of transformation driven by PTEN loss (Fig. 2J). Although the IFN pathway does not appear to be required for the process of transformation *in vitro*, the cells emerged from the stress of growth factor deprivation would be forced to adapt to any unfavorable consequences of the activated IFN signaling, which is often associated with cytotoxicity.

### PTEN-deficient tumors exhibit robust, tumor cell-intrinsic interferon signaling

We next asked whether the epithelia cell-intrinsic interferon response is also activated in human tumors with a PTEN-low status. To circumvent confounding differences in the immune composition of tumor samples, we examined scRNA-Seq data generated from human breast tumors^34,35^ (Fig. 3A top, S6). Tumor epithelial cells in triple-negative breast cancer (TNBC) – in comparison to their counterparts in HER2+ or ER+ tumors – displayed low *PTEN* expression that is associated with high expression of *STAT1* and other interferon-stimulated genes (ISGs) (Fig. 3A bottom right, S6). Notably, myeloid cells from TNBC were not apparently distinct from those found in HER2+ breast cancer in terms of ISG expression (Fig. 3A bottom middle), emphasizing a cancer cell-intrinsic activation of interferon signaling in PTEN-low TNBC tumors.

**Figure 3.**
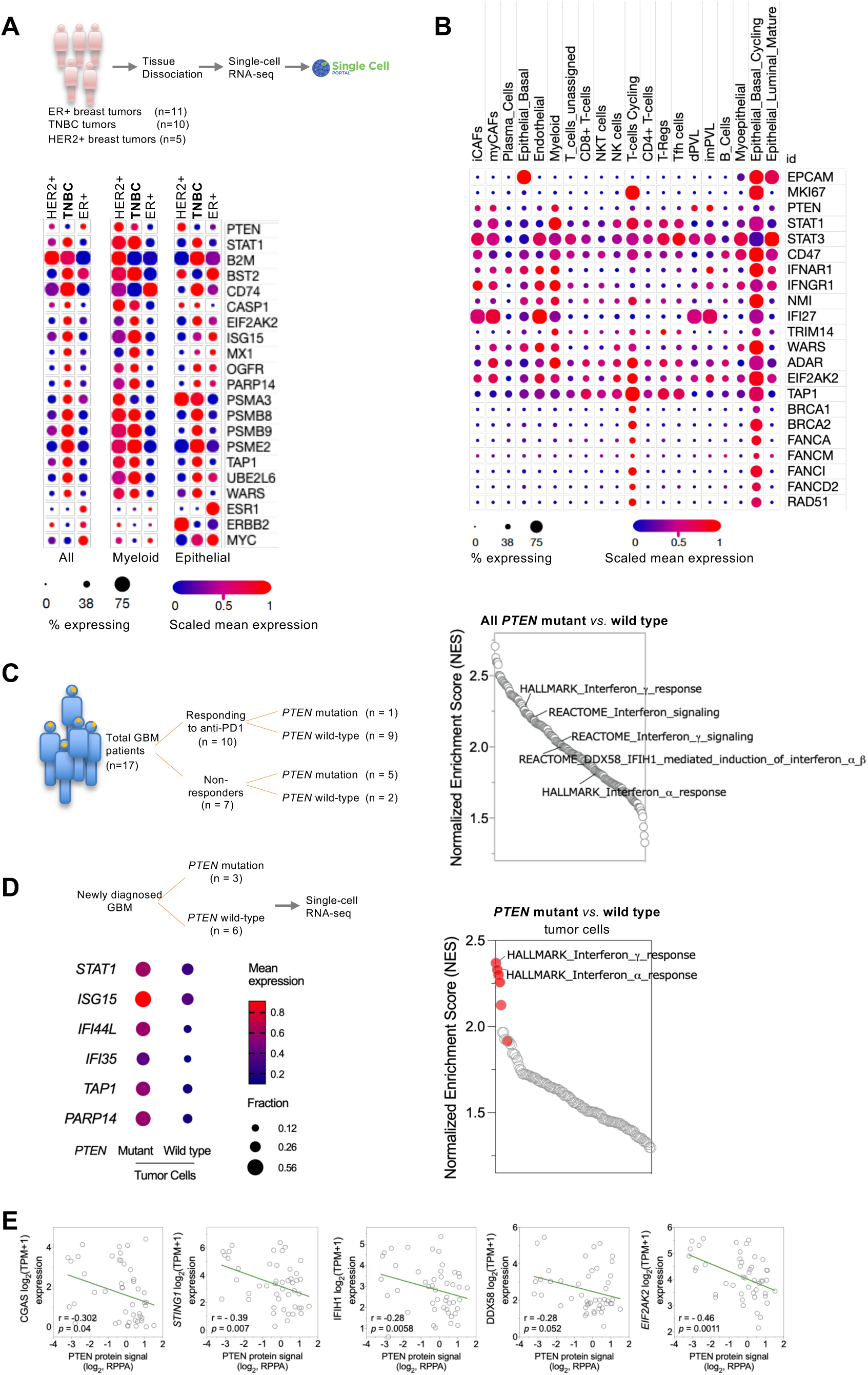
Tumor cells with low expression of PTEN exhibit robust activation of the interferon-STAT pathway. (A) (Top) Schematic of single-cell RNA-seq data generation and analysis. (Bottom) Heatmap of indicated gene expression among all cells in the indicated types of tumors (Left), myeloid cells (Middle), or epithelial cells (Right). (B) Dot plot depicting the expression of the indicated genes among twenty cell types present in triple-negative breast tumors. (C) (Left) Schematic of bulk RNA-seq for glioblastoma samples isolated from patients prior to immune checkpoint blockade therapy. (Right) Gene expression of all PTEN-mutated tumors was compared to that of all PTEN wild type tumors, and the results were subjected to gene set enrichment analysis. Gene sets related to interferon signaling are indicated. (D) (Top left) Breakdown of single-cell RNA-seq for glioblastoma samples isolated from newly diagnosed patients. (Bottom left) Tumor cells were identified as described in the methods. For the ISGs shown, the mean expression and fraction of cells expressing them in PTEN-mutated or wild type samples were plotted. (Right) Gene expression of PTEN-mutated tumor cells was compared to that of PTEN wild type tumor cells, and the results were subjected to gene set enrichment analysis. Gene sets related to interferon signaling are noted in red. (E) PTEN protein abundance is significantly correlated with the expression of interferon-stimulated genes (ISGs) that encode nucleic acid sensors. Gene expression (determined by RNA-seq) and PTEN protein abundance (quantified by reverse-phase protein array) in human breast cancer cell lines (n = 47) were downloaded from DepMap and analyzed in GraphPad Prism.

We further investigated ISG expression in TNBC cells, and found that among all tumor epithelial cells, cycling epithelial cells – identified by the expression of the cell proliferation marker *MKI67* and the epithelial marker *EPCAM*, have little to no PTEN expression, but display robust ISG expression (Fig. 3B, S7A). Furthermore, this elevated expression of ISGs in cycling epithelial cells was even greater than that found in myeloid or cycling T cells present in the tumor microenvironment (Fig. 3B, S7B). Thus, PTEN-low TNBC tumor cells display a high level of interferon signaling gene expression specifically in the cycling tumor epithelial cells.

The activation of interferon signaling also occurs in other types of tumors with a PTEN-low status. In a cohort of glioblastoma (GBM) patients, PTEN mutation is associated with resistance to immune-checkpoint blockade therapy^17^(Fig. 3C left). We compared the bulk gene expression in all PTEN mutated tumors with that in wild type tumors regardless of their response to the treatment, and found that PTEN mutated tumors have significantly enriched expression of interferon signature genes (Fig. 3C right, Fig. S8A). Notably, we did not observe a significant difference between the mRNA levels of many immune cell-specific markers, including those of macrophages and T cells, two major types of immune cells present in glioblastoma (Fig. S8B-C). Therefore, the tumor cells are likely a major contributor to the differences in ISG expression between PTEN mutant and wild type tumors. We next analyzed scRNA-Seq data in a cohort of newly diagnosed GBM^36^, and found significantly higher expression of ISGs in PTEN mutated tumor cells, compared to their counterparts with wild type PTEN (Fig. 3D, S8D-E). Furthermore, the expression of these genes in tumor cells is comparable to that of the immune cells, including monocytes and T cells in the tumor microenvironment (Fig. S8D, 8F).

Given the complexity of human cancer genetics, we next asked if Pten inactivation in genetically engineered mouse models (GEMs) also results in activation of interferon signaling in tumor cells. In an ovarian tumor model driven by *Pten* loss together with three other oncogenic alterations^37^, cancer cells express interferon response genes as robustly as monocytes found in the ovarian tumor microenvironment (Fig. S9A). Such profound activation of interferon signaling is also observed in tumor cells from a mouse model generated by ablating *Pten* in prostate luminal epithelial cells^38^. These Pten-deficient tumor cells, similar to one sub-population of monocytes in the tumor microenvironment, strongly express ISGs, including *Stat1*, *Stat2*, *Irf9*, *Sting*, and *Pkr* (Fig. S9B).

Finally, in established human cancer cell lines, we also observed a significant reverse correlation between PTEN protein abundance and the transcript levels of ISGs, such as those encoding double-stranded DNA or RNA sensors (Fig. 3E; Fig. S9C-D). This association further reinforces the observation of cancer cell-intrinsic interferon signaling in PTEN-low cancer cells.

### PTEN low status renders cancer cells resistant to immunosurveillance but causes a synthetic sensitivity to CDK12/13 inhibition

We next investigated whether PTEN status determines cancer cell-intrinsic interferon response. Given the crucial role of IFN-γ pathway in tumor surveillance^39–43^, one pressing question is whether the elevated interferon response in cancer cells may force cancer cells to adapt to the cytotoxic effects of interferons and consequently gain resistance to immune clearance.

To study if PTEN status may have a causal role for the induction of IFN signaling, we examined the expression of ISGs in PTEN-deficient cancer cells upon ectopic expression of PTEN. PTEN expression reduced the protein abundance of several key ISG-encoded proteins critical for IFN response (Fig. 4A). We also treated the cells with IFN-γ to measure their response to its cytotoxicity. Intriguingly, cells with PTEN re-expressed became profoundly sensitive to the killing induced by IFN-γ (Fig. 4B-C). Moreover, in co-culture models comprising tumors cells and NK cells or cancer cell-recognizing T cells, the ectopic expression of PTEN rendered cancer cells significantly sensitive to immune cell-mediated killing (Fig. 4D, S10A).

**Figure 4.**
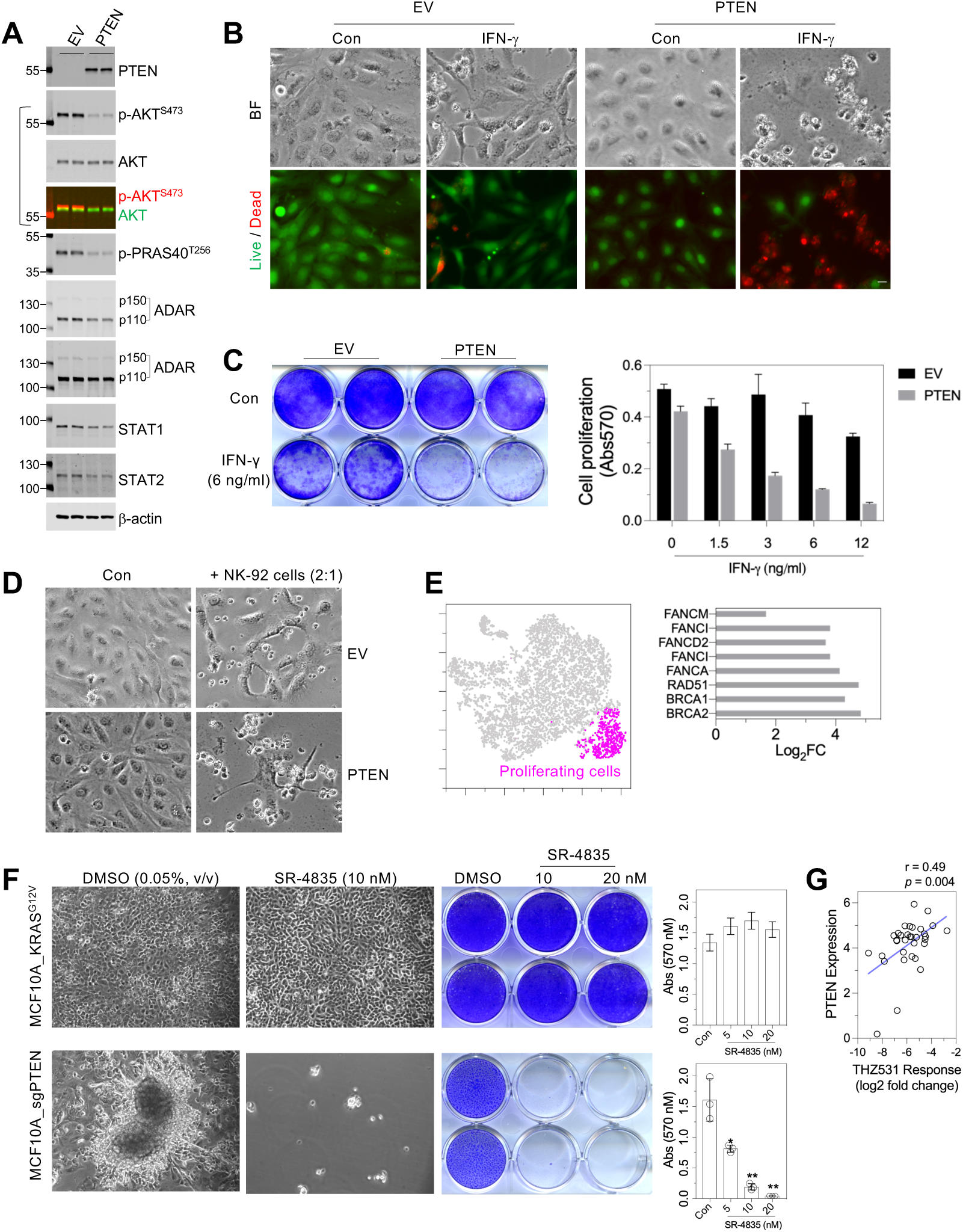
PTEN-deficient cells are resistant to IFN-γ and cytotoxic immune cells, but highly sensitive to CDK12/13 inhibition. (A) Fluorescent immunoblotting of whole-cell lysates from BT-549 cells stably transduced with an empty vector (EV) or a lentivirual vector encoding PTEN (PTEN). Merged blots for phosphorylated and total AKT are shown. A long-exposure blot for ADAR is shown to visualize the p150 isoform. (B) BT-549 cells with an empty vector (EV) or PTEN addback were seeded in 12-well plates with or without IFN-γ (6 ng/ml). Seven days post-treatment, cells were subjected to live/dead staining. Bright-field and fluorescent images were captured with a 10x objective lens. (C) Cells were treated as in (B), fixed, and stained with crystal violet (Left), followed by quantification of staining (Right; mean ± SD). (D) Co-culture of cancer cells with NK-92 cells. BT-549 cells with an empty vector or PTEN addback were left untreated or co-cultured with NI-92 cells (effector-to-target ratio: 2:1). After one week of culture, bright-field images were captured. (E) Expression of the indicated DNA damage response (DDR) genes (Right) in the cluster of proliferative cells (Left) within MCF10A_sgPTEN transformed colonies. Data are derived from single-cell RNA-seq. (F) MCF10A cells stably transduced with KRAS^G12V^ or sgPTEN were subjected to serum, EGF, and insulin starvation in the presence of vehicle or SR4835. After four weeks, cells were imaged (Left), fixed, and stained with crystal violet (Middle), followed by staining extraction and quantification (Right; mean ± SD; * p < 0.05, ** p < 0.01). (G) Correlation analysis of PTEN expression and sensitivity to THZ531 in breast cancer cell lines (n = 33). Analysis was performed using https://depmap.org/portal/, with data downloaded and plotted in GraphPad Prism.

These data suggest that restoration of PTEN in PTEN-deficient cancer cells can re-sensitize them to immune recognition, however, this approach as a potential therapeutic strategy has its inherent technical challenges. We thus considered whether PTEN loss-driven transformation may generate any synthetic dependence that could be readily exploited for therapeutic intervention. We further analyzed single-cell gene expression in the *in vitro* transformation model and found that Cluster 6, the population of proliferating cells that is the mostly separated from the rest of cells, demonstrate robustly elevated expression of genes encoding DNA damage response (DDR) proteins (Fig. 4E). Notably, robust expression of DDR genes was also found in PTEN-low and cycling epithelial cells in human TNBC samples (Fig. 3B), suggesting that activation of DNA damage response may be common characteristic of PTEN-low cycling tumor cells.

Reminiscent of activated DNA damage response documented in early stages of human cancer development^44,45^, these observations suggest that PTEN-deficient proliferating tumor cells may have an elevated DNA damage response leading to induction of DDR gene expression for adaptation. The observations also led us to hypothesize that PTEN-deficient cancer cells may be dependent on a high level of DDR gene expression, and particularly on one DDR protein CDK12 – a transcriptional cyclin-dependent kinase that is known to be critical for the transcription of DDR genes^46–49^. We treated PTEN-deficient transformed cells with various kinase inhibitors that target transcriptional CDKs or PIK3CB/p110β, and found that THZ531, a covalent inhibitor selectively suppressing CDK12 as well as its paralog protein CDK13^50^, nearly eliminated the transformed colonies (Fig. S10B-C). The effect of CDK12/13 inhibition in established sgPTEN colonies was far more profound than those arising from CDK7, CDK8, or PIK3CB/p110β inhibition (Fig. S10B). We further tested a second structurally independent CDK12/13 inhibitor SR-4835^51^, and found that the compound was able to eliminate PTEN-deficient transformed cells at low nanomolar concentrations (Fig. S10D). Notably, similar doses of SR-4835 was well-tolerated in untransformed cells (Fig. S10E). We also studied whether such an exquisite sensitivity is limited to cells transformed by PTEN loss. To this end, we introduce a constitutively active form of KRAS (KRAS^G12V^) into MCF-10A cells and found that these cells were able to survive after prolonged starvation of serum and growth factors. However, and distinct from cells with PTEN knock out, cells expressing KRAS^G12V^ did not grow into colonies and rather presented an apparent visual homogeneity. Interestingly, KRAS KRAS^G12V^-transformed cells were completely immune to CDK12/13 inhibition, suggesting that the extreme sensitivity to CDK12/13 inhibition is specific to cells transformed by PTEN loss (Fig. 4F). Consistent with these observation in epithelial cells, a robust correlation was also found between PTEN expression and the cytotoxicity of the CDK12/13 inhibitor THZ531 among breast cancer cell lines, *i.e.*, the less PTEN cancer cells express, the more sensitive they are to CDK12/13 inhibition (Fig. 4G).

## Discussion

Integrating multi-omics analyses and a PTEN loss-driven transformation model, we have found that human PTEN-low cancer cells uniquely depend on PIK3CB/ p110β, a characteristic that breast epithelial cells can acquire during PTEN loss-induced cell transformation. The transformed cells demonstrate a gene expression landscape that closely resembles human basal-like breast cancer, with a striking degree of heterogeneity in both morphology and pattern of gene expression. Surprisingly, this *in vitro* transformation model, arising from untransformed epithelial cells, reveals a striking activation of an interferon response, as indicated by hyperactivated gene expression of interferon-stimulated genes including both core transcription factor STAT1 and those involved in antigen processing and presentation. Such epithelial cell-intrinsic activation of IFN signaling in the absence of PTEN expression is also found in PTEN-low/deficient human breast cancer and glioblastoma, as well as mouse tumors generated via genetic engineering to ablate the *Pten* gene. Thus, the tumor cell intrinsic activation of interferon response appears to be an inherent property of tumor cells driven by loss of the tumor suppressor PTEN.

What is the functional relevance of tumor cell-intrinsic activation of interferon signaling in PTEN-low tumors? In the tumor microenvironment, interferon-γ (IFN-γ) is secreted by activated cytotoxic T cells and acts on tumor cells to induce cell death and/or growth inhibition, as well as to upregulate the expression of proteins involved in antigen presentation. The IFN-γ pathway was found to be critical for tumor surveillance *in vivo*^39–41^, for chimeric antigen receptor (CAR) T cell cytotoxicity towards solid tumor cells^52^, and for T cell-mediated killing of tumors cells *in vitro* in co-culture models consisting of mouse tumor cells and tumor cell-recognizing T cells^42,43^. In addition, IFN-γ signaling is found to drive clinical response to ICB in melanoma patients^53^, and its activation is strongly associated with improved overall survival^54^. However, it is not exactly clear whether the killing effect arises from IFN-γ itself, IFN-γ-induced antigen presentation, or both. Notably, loss-of-function mutations in the JAK1 or JAK2, kinases essential to the IFN-γ pathway, were observed in melanoma patients who had acquired resistance to PD-1 blockade therapy^55^, as well as in patients with primary resistance to anti-PD-1 treatment^56^. In addition, genomic defects in IFN-γ pathway genes were observed in melanoma patients who had failed to respond to anti-CTLA-4 therapy^57^. Importantly, ablating JAK1 or JAK2 in a human melanoma cell line impairs IFN-γ-induced tumor cell growth arrest, but does not impact tumor antigen presentation^55^. A dominant role of IFN-γ-mediated tumor cell killing was recently discovered by studies that develop CD4+ CAR T cells for cancer therapy; IFN-γ, instead of perforin-mediated pathway, is found to be responsible for tumor elimination, and this killing action can occur in distant tumor cells via diffusion of IFN-γ□secreted by CD4+ CAR T-cells^58,59^.

Together, these studies implicate enhancing IFN-γ-elicited killing of tumor cells as a major effect of immune checkpoint blockade therapy. Therefore, persisting tumor cell-intrinsic activation of the IFN pathway, which we observe in PTEN-low tumors, suggests that the tumors cells might have been forced to adapt to this suppressive mechanism during the earlier process of tumorigenesis, and would be resistant to treatments designed to augment this pathway.

In support of this IFN adaptation model, we find that restoring PTEN expression in PTEN-deficient cancer cells strikingly re-sensitizes cells to the cytotoxicity of IFN-γ, as well as to both cytotoxic NK cells and T cells. Notably, a previous study found that reducing PTEN expression in a PTEN-proficient human melanoma cell line reduces T cell-mediated anti-tumor activity^15^. Together these observations suggest that re-expressing PTEN in PTEN-deficient tumors could be a strategy to force tumor cells respond to immune checkpoint therapies, which have extremely limited efficacy in human tumors with a loss of PTEN expression/function^15–19^. Interestingly, a recent study demonstrates that restoring PTEN expression in tumor cells via nanoparticle-delivered mRNA enhances response of allografted mouse tumors to immune checkpoint blockade^60^. Despite these advances, the mechanisms by which PTEN status regulates cancer cell response to cytotoxic immune cells need to be further elaborated. Previous studies demonstrate that PTEN loss can lead to the upregulation of immunosuppressive cytokines such as CCL2 and VEGF^15^, as well as STAT3 activation^21^ (a transcription factor that can suppress the expression of immune stimulatory cytokines). Distinct from these investigations, our study points to STAT1 as a key factor that is strongly induced during transformation induced by PTEN loss. Previous studies have found that the alpha form of the STAT1 protein harbors a serine residue (Ser727) whose phosphorylation is known to be important for the maximal activation of transcription by STAT1^61^. Notably, we found that Ser727 phosphorylation of STAT1 is susceptible to the inactivation of PI3K/AKT pathway (*data not shown*), suggesting a potential link between PTEN status and IFN signaling.

The IFN adaptation model we have proposed for PTEN-low tumor largely relies on the effect of IFN-mediated cytotoxicity on tumor cells. However, IFN response also induces the expression of certain ISGs (*e.g.*, PD-L1) that leads to T cell inhibition. Indeed, recent studies demonstrate that persistent IFN signaling in tumors cells causes adaptative resistance to ICB^62,63^, and delayed administration of JAK inhibitors can improve response to ICB in both mouse tumor models^62,63^ and cancer patients^64,65^. Therefore, PTEN-low tumors may not only adapt to the cytotoxic effects of IFN, but can also achieve adaptative resistance to ICB through induction of both IFN response-dependent and -independent pathways to achieve T cell inactivation and exhaustion^15,21^. These two types of mechanisms, adaptation to IFN cytotoxicity and immunosuppression, likely co-exist in PTEN-low tumors. Their contribution to the outcome of tumor growth and/or response to ICB may depend on whether tumors cells have naïve sensitivity to IFN cytotoxicity and thus may vary with the genetic backgrounds of particular tumors and cell types.

In summary, our study reveals a robust interferon response in epithelial cells transformed by PTEN loss *in vitro*, and in PTEN-low/null tumor cells from clinical samples including those of triple-negative breast cancer and glioblastoma. Our experimental results and analyses of human data support an IFN cytotoxcity adaption model, wherein PTEN-low/null tumor cells are forced to adapt to the cytotoxicity associated with interferon responses and, therefore, are rendered resistant to therapies that rely on the activation of interferon pathway to eliminate the tumor cells. Future investigation is warranted to study whether PIK3CB/p110β□inhibition mimics the restoration of PTEN expression to sensitize tumor cells to immune checkpoint blockade therapies, as well as the mechanistic actions of CDK12/13 – inhibition of which was identified as an effective new therapeutic strategy for PTEN loss-driven tumors.

## ACKNOWLEDGMENTS

We are grateful to Drs. Justin Becker, Bradley Bernstein, Gordon Freeman, Lewis Cantley, Tao Zou (Dana-Farber Cancer Institute), Yang Shi (Ludwig Oxford), and Brian Schaffhausen (Tufts Medical School) for fruitful discussion. We thank Dr. Robert Weinberg (MIT) for sharing the mouse ovarian cancer cell model PPNM. We also thank Drs. Jing Ni, Xin Chen, Qiwei Wang, and Johann S. Bergholz for discussion and sharing reagents. Y.W. acknowledges funding by the Eleanor and Miles Shore Faculty Development Awards Program of Harvard Medical School. J.J.Z acknowledges funding by NIH grant R35 CA210057. T.M.R. acknowledges funding by NIH grant R35 CA231945.

## AUTHOR CONTRIBUTIONS

Conceptualization: Y.W., P.B. and T.M.R. Formal analysis: Y.W., C.M., J.C. and D.L. Investigation: Y.W., C.M. and H.G. Funding acquisition: J.J.Z. and T.M.R. Methodology: Y.W., C.M. and D.L. Project administration: T.M.R. Supervision: Y.W., D.L. and T.M.R. Writing - original draft: Y.W. and T.M.R. Writing - review & editing: Y.W., D.L., P.B., J.J.Z. and T.M.R

## DECLARATION OF INTERESTS

T.M.R. is a Scientific Advisory Board member for Shiftbio and K2B Therapeutics and is a co-founder of Geode Therapeutics Inc. J.J.Z. is a co-founder and board director of Crimson Biotech Inc. and Geode Therapeutics Inc. D.L. received speaking honorariums and travel fees from Genentech and consulting fees from Oncovalent Therapeutics, not pertinent to or affected by the content of this publication. All other authors declare no competing interests.

---

**Figure S1.**
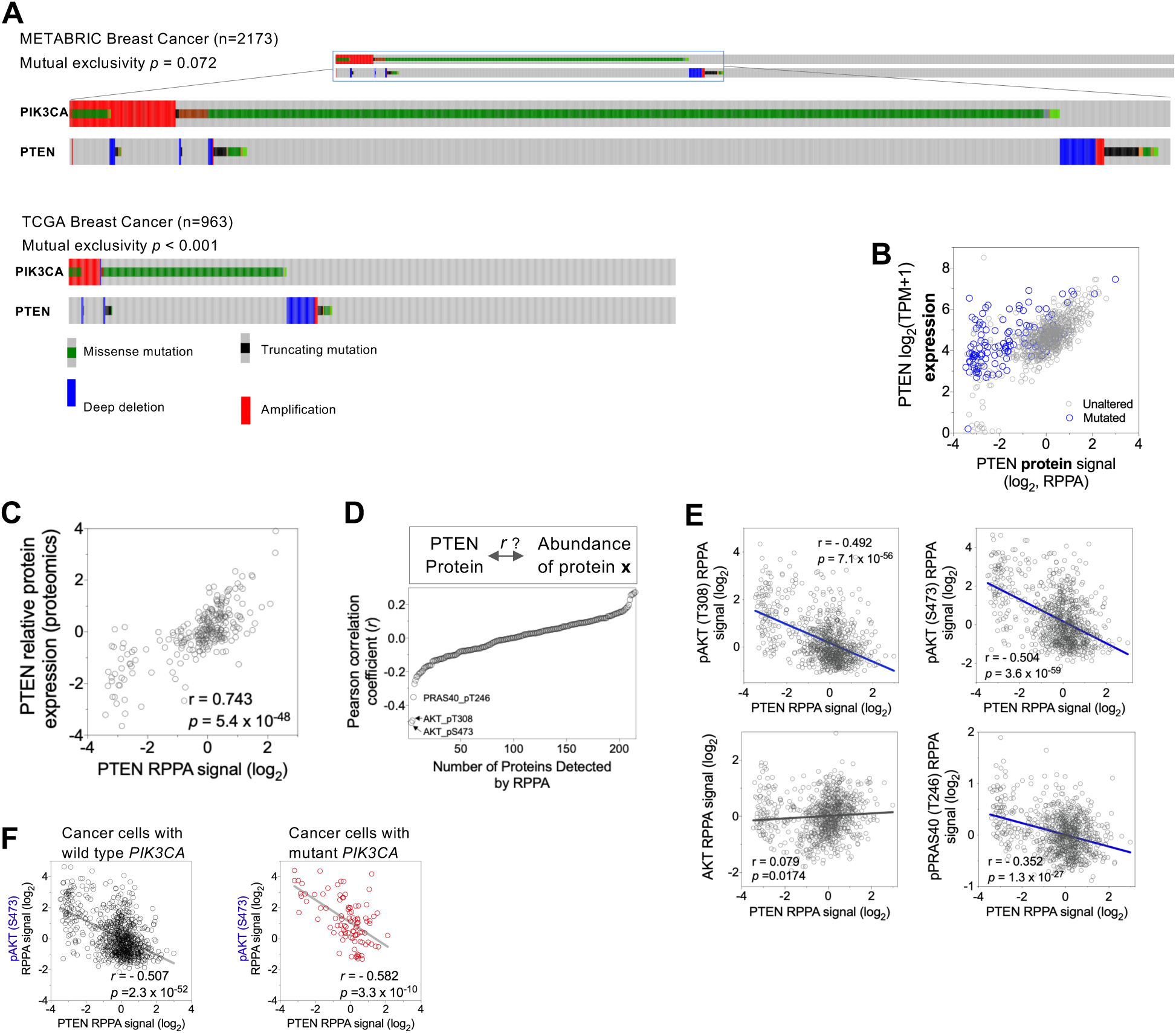
PTEN alterations in human cancer, and the use of PTEN protein abundance as a molecular marker to accurately reflect PTEN status. (A) Status of PIK3CA and PTEN genetic alterations in human breast cancer. The OncoPrint visualization was derived from TCGA breast cancer (TCGA, 2012) and METABRIC breast cancer cohorts (Curtis et al., 2012), available at the cBioPortal for Cancer Genomics (www.cbioportal.org; Cerami et al., 2012; Gao et al., 2013). Note that PTEN gene deletion or mutation is mutually exclusive with alterations of PIK3CA. (B) A plot illustrating PTEN protein abundance (x-axis) and its transcript level (y-axis) in cancer cell lines (n = 961). Note that for a significant number of cell lines, PTEN protein abundance is decoupled from PTEN genetic status (e.g., many cell lines with wild type PTEN genes, as depicted by gray circles in the bottom left, have low levels of expression), and cancer cells harboring PTEN mutation have similar mRNA expression to those with wild type PTEN. (C) Correlation analysis of PTEN protein abundance assayed by RPPA (x-axis) versus the abundance assayed by proteomics (y-axis). Pearson correlation coefficient (r) and p-value are indicated. (D) PTEN protein abundance determined by RPPA is analyzed for correlation with other proteins in the dataset. Note that pAKT_S473, pAKT_T308, and pPRAS40_T246 are the top three hits for the analysis. (E) Correlation analysis of PTEN protein abundance assayed by RPPA (x-axis) versus the indicated (post-translationally modified) proteins. Pearson correlation coefficient (r) and p-values are indicated. (F) Correlation analysis of PTEN protein abundance assayed by RPPA (x-axis) versus pAKT_S473 signal in cancer cell lines with wild type (left) or mutated (right) *PIK3CA*. Note that the correlation is significant, irrespective of *PIK3CA* status.

**Figure S2.**
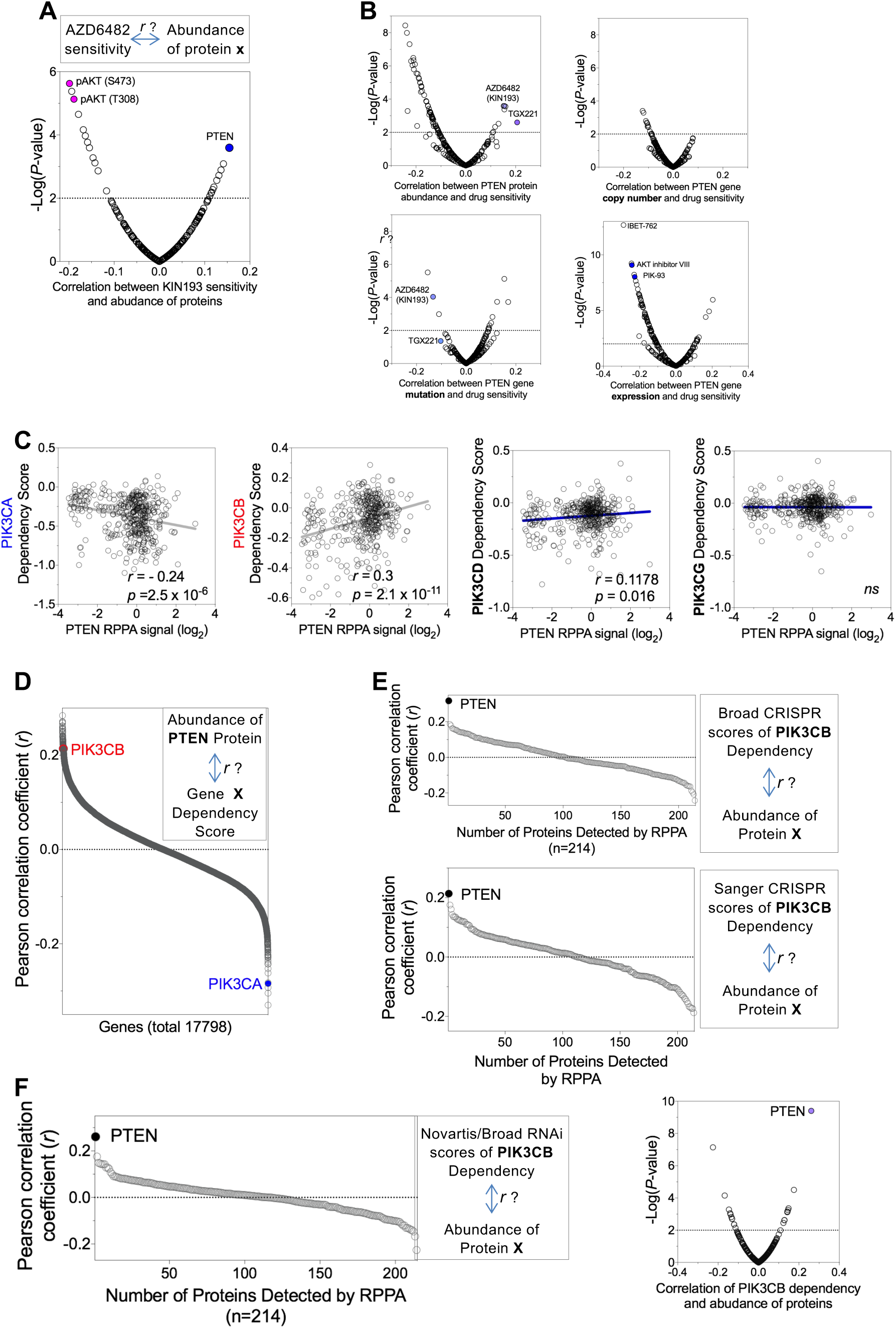
PTEN-low cancer cells prominently depend on PIK3CB. (A) AZD6482 (also named KIN193) sensitivity scores were used as bait for correlation analysis with protein abundance data in the RPPA dataset. PTEN protein was the greatest positive correlate, while phosphorylated AKT was the greatest negative correlate. (B) Cancer cell drug sensitivity data generated by the GDSC project was analyzed for correlation with PTEN protein (Top left), PTEN gene copy number (Top right), PTEN gene mutation (Bottom left), or PTEN gene expression (bottom right). Note that compared with the other molecular traits of PTEN, PTEN protein abundance has the strongest association with response to PIK3CB inhibitors TGX221 and AZD6482/KIN193. The top left plot is similar to Figure 1A and has less compounds (n = 266) included. (C) Correlation analysis between the PTEN protein abundances (x-axis) and dependency score of indicated genes (PIK3CA/p110α, PIK3CB/p110β, PIK3CD/p110δ, PIK3CG/p110γ). Dependency scores are from the Avana library CRISPR screen (Doench et al., 2016). Pearson correlation coefficient (r) and p-value are indicated; *ns* denotes “not significant”. (D) Correlation analysis between PTEN protein abundance and genome-wide dependency scores from the Sanger CRISPR screen (Behan et al., 2019). Pearson correlation coefficient (y-axis) of each individual gene is plotted. (E) PIK3CB dependency scores from Broad Avana CRISPR screen (Top) or Sanger screen (Bottom) were analyzed for correlation with the RPPA dataset (total 214 proteins, including total and/or post-translationally modified proteins). Note that PTEN is the top hit for both analyses. (F) Analysis was performed as in (E) using PIK3CB dependency scores from the Novartis/Broad RNAi screens (McDonald et al., 2017). The volcano plot on the right shows that only a handful of proteins demonstrate statistically significant correlation with PIK3CB dependence.

**Figure S3.**
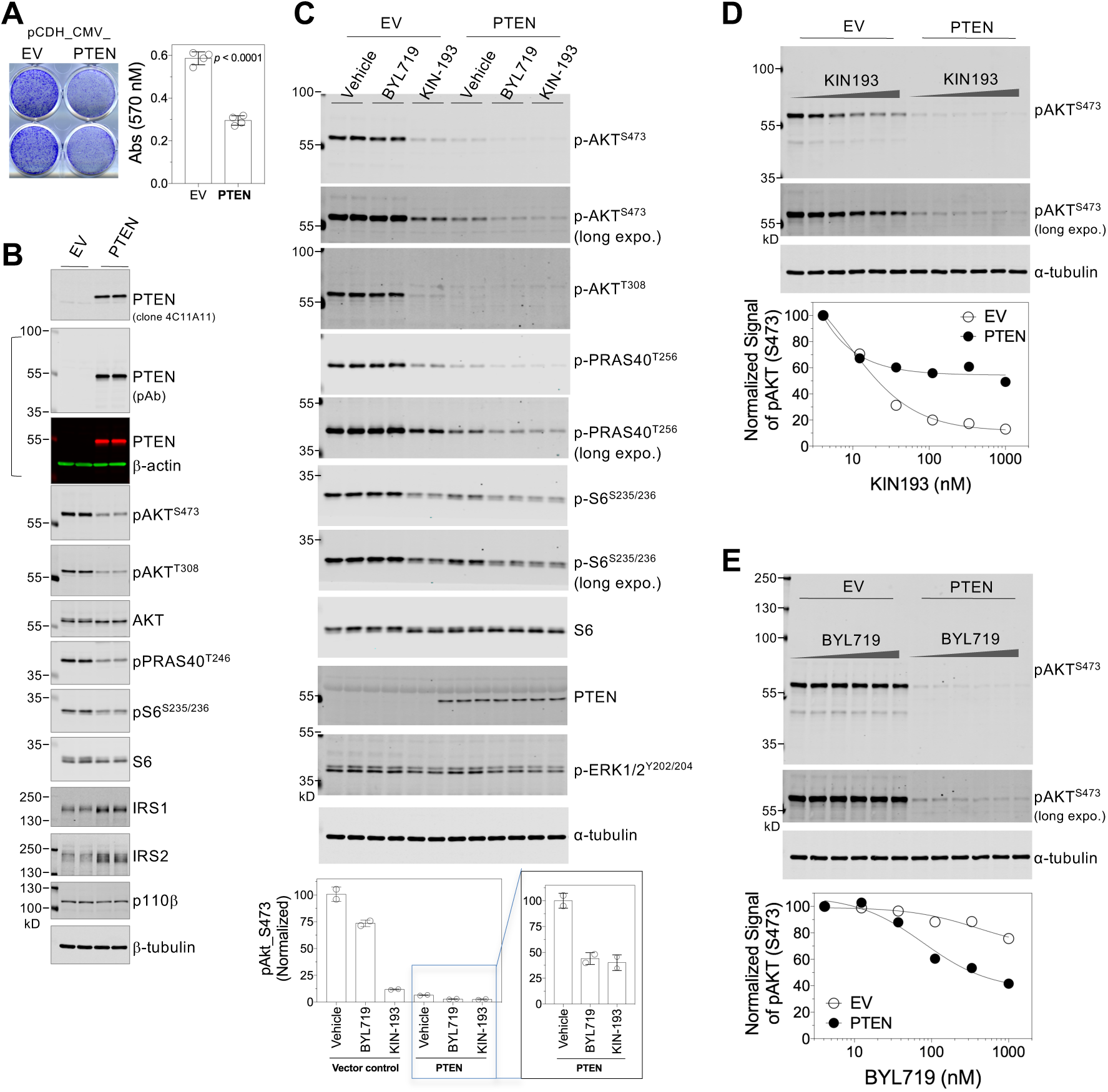
Ectopic expression of PTEN in PTEN-deficient cells alters the cellular response to PI3K p110α and p110β inhibition. (A) Triple-negative breast cancer line MDA-MB-468 cells transduced with empty vector (EV) or an untagged form of PTEN were seeded in 12-well plates at the density of 10,000 cells *per* well, and in 7 days fixed and stained with crystal violet (left). The staining was then dissolved for quantification of the absorbance (right). Note that forced expression of PTEN impairs cell growth, and a significant amount of time was required before cells achieve adaptation and a sufficient number of cells become available for assays. (B) Cells as in (A) were subjected to lysis with 1x SDS sample buffer followed by fluorescent immunoblotting using the indicated antibodies. A merged image is shown for PTEN (derived from the use of a rabbit polyclonal antibody) and β-actin (from the use of a mouse monoclonal antibody) as the loading control. (C-E) Cells were treated with vehicle control (0.1% DMSO, v/v), PIK3CA/p110α inhibitor BYL719 (also named alpelisib), or PIK3CB/p110β inhibitor KIN193 (also named AZD6482) for 1 hour in complete medium, before being lysed for fluorescent immunoblotting. The fluorescent signal of pAKT^S473^ was normalized to that of α-tubulin (bottom graph in C-E). Non-linear regression was performed for dose response of pAKT^S473^ signal (D-E; GraphPad Prism 7) Note that for all immunoblots, cells were lysed directly with 1x SDS sample buffer, followed by fluorescent immunoblotting using the indicated antibodies. The protein marker shown at the very left of each blot (PageRuler Plus Prestained Protein Ladder, Thermo Fisher Scientific) has near-infrared fluorescence, a signal that can be detected using the 700-nm channel in the Odyssey CLx Infrared Imaging System.

**Figure S4.**
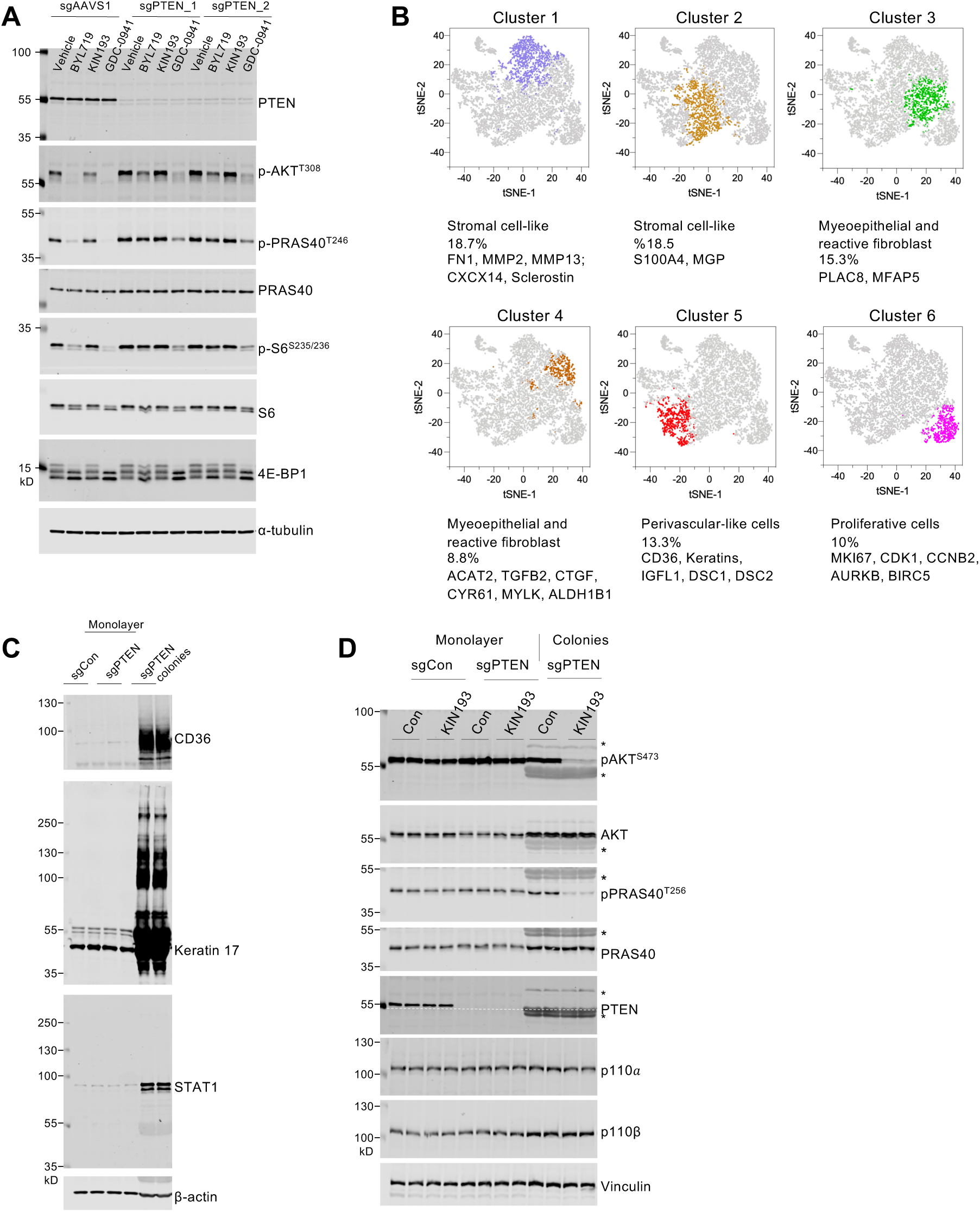
PTEN loss renders transformed cells sensitive to PIK3CB inhibition. (A) CRISPR/Cas9-mediated gene editing of PTEN does not immediately confer cells sensitivity to PI3KCB inhibition. Human mammary epithelial cells (HMEC) stably transduced with a control or PTEN-targeting guide were seeded and treated the next day with PIK3CA inhibitor BYL-719, PIK3CB inhibitor KIN193/AZD6482, or pan-PI3K inhibitor GDC-0941 (all 1 μM) in complete medium for 1 hour. Cells were lysed with 1x SDS sample buffer and subjected to fluorescent immunoblotting. (B) Transformed MCF10A_sgPTEN colonies were subjected to single-cell RNA-seq, and t-SNE projections of cells colored by automatic clustering is shown. Clusters of cells and their respective markers are indicated. (C) Fluorescent immunoblotting of whole-cell lysates from MCF10A_sgCon cells, MCF10A_sgPTEN monolayer cells, and MCF10A_sgPTEN transformed colonies. The molecular weight of fluorescent protein markers is indicated. Note the strong induction of proteins in colonies, including CD36, Keratin 17, and STAT1. (D) Monolayer cells (sgCon, sgPTEN) or sgPTEN colonies were treated with control (0.1% DMSO, v/v) or 1 μM KIN193/AZD6482 for 1 hour, followed by lysis with sample buffer and immunoblotting. Note that non-specific signals from the colony samples in all blots are labeled with an asterisk (*).

**Figure S5.**
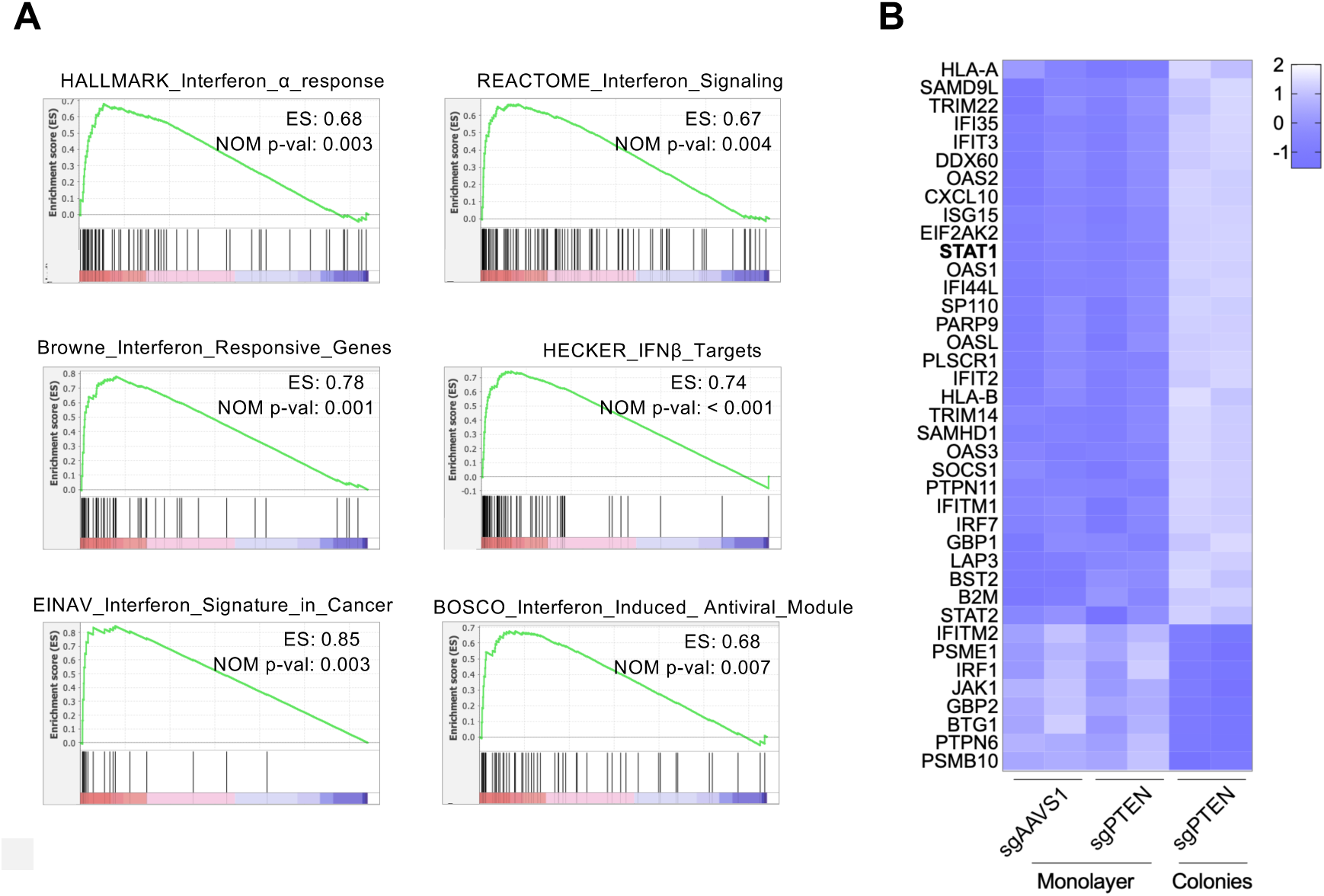
PTEN loss-transformed cells exhibit a profound activation of the interferon response. (A) GSEA plots of the indicated interferon signatures for genes significantly altered in transformed MCF10A_sgPTEN colonies compared to their pre-transformed counterparts. (B) Heatmap of selected genes from interferon response signatures in the indicated samples. Note that sgPTEN colonies (transformed cells) exhibit increased expression of many interferon-stimulated genes (ISGs), including STAT1 and STAT2.

**Figure S6.**
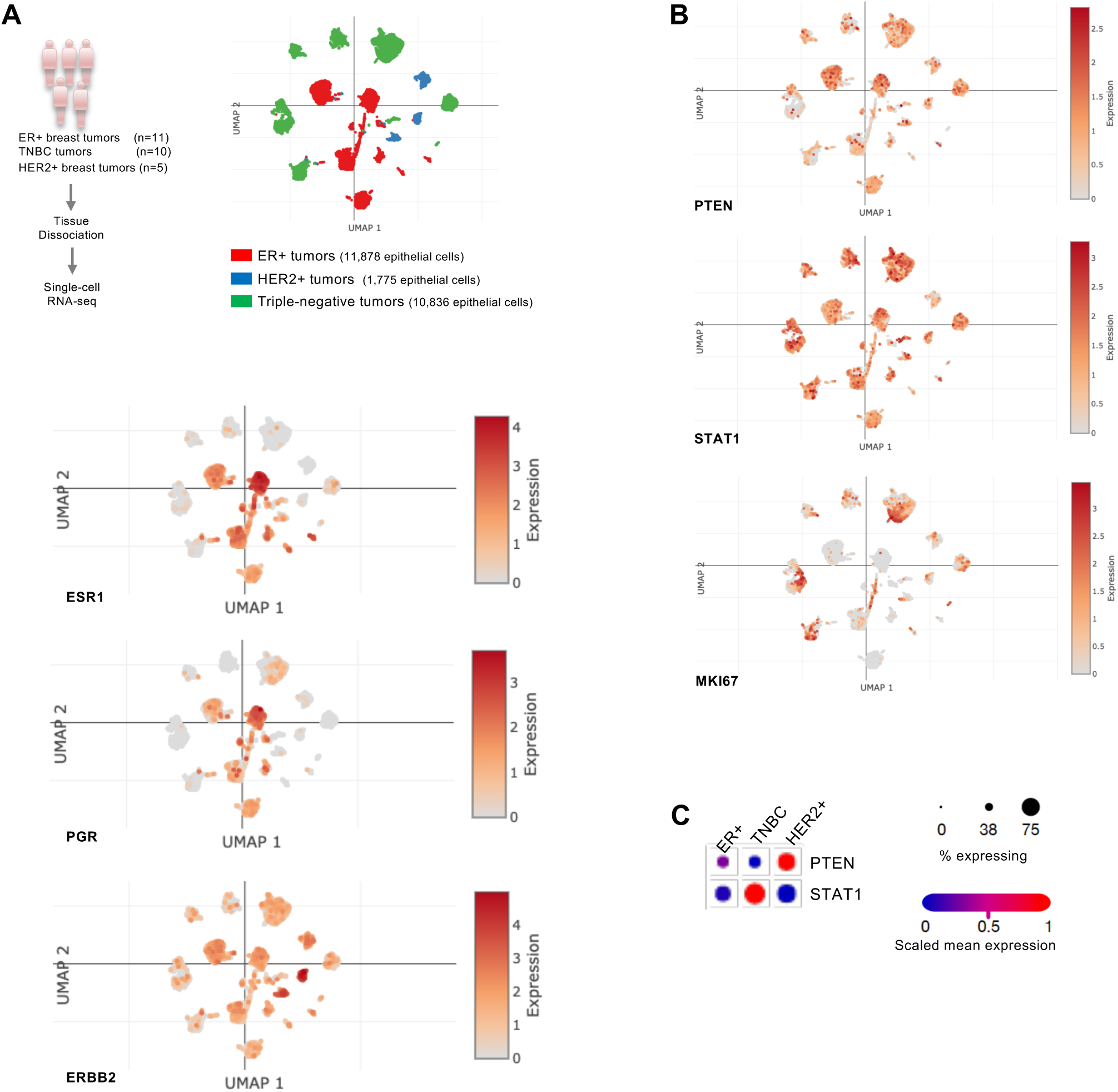
Tumor cells in triple-negative breast cancer exhibit low PTEN expression and robust, cancer cell-intrinsic expression of STAT1. (A) (Top left) Schematic of single-cell RNA-seq data generation and analysis (Wu et al., 2021). UMAP (Uniform Manifold Approximation and Projection) visualization of epithelial cells from 26 primary breast tumors (top right), and (bottom) the expression of clinical markers for breast cancer subtypes, including estrogen receptor (ESR1), progesterone receptor (PGR), and human epidermal receptor tyrosine kinase HER2 (ERBB2). (B) Normalized expression of the indicated genes in tumor epithelial cells. Note that triple-negative breast cancer cells have low expression of PTEN and high levels of STAT1 and MKI67. (C) Summary of PTEN and STAT1 expression data from panel (B).

**Figure S7.**
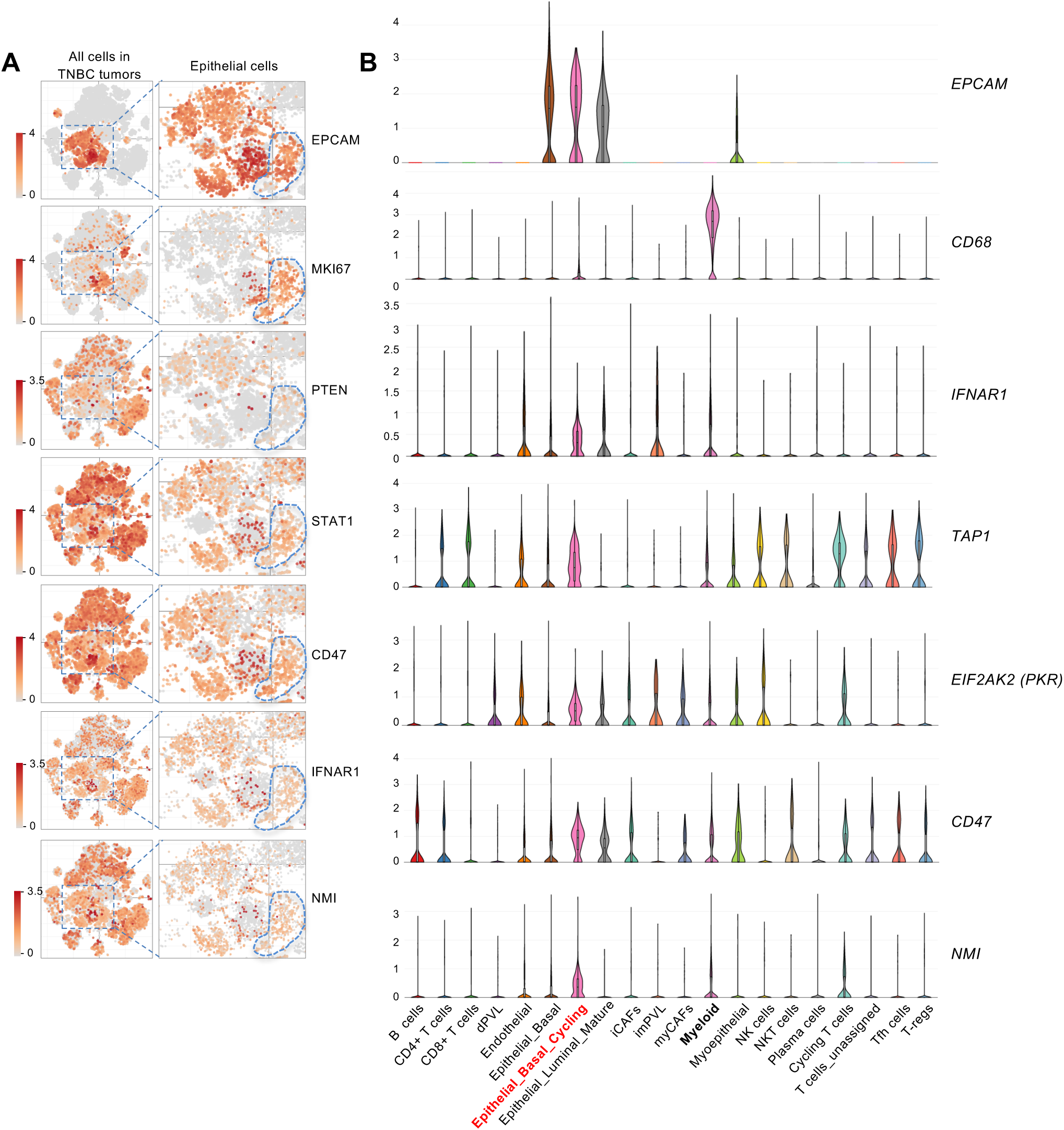
Robust expression of interferon-stimulated genes in cycling tumor cells of triple-negative breast cancer. Triple-negative breast tumors from ten patients were subjected to tumor dissociation and single-cell RNA-seq (Wu et al., 2020). Data analysis was performed using the Broad Institute’s Single-Cell Portal (https://singlecell.broadinstitute.org/single_cell). (A) t-SNE plots of expression for the indicated genes, with the plots on the right showing zoomed-in views of epithelial cells. Note that cycling breast tumor epithelial cells (dotted circle), marked by EPCAM and MKI67, have low expression of PTEN and higher expression of the indicated interferon-stimulated genes (ISGs) compared to non-cycling tumor cells. (B) Violin plots of gene expression for the indicated cell types. Note that cycling tumor cells (annotated as “Epithelial_Basal_Cycling”) demonstrate robust expression of the indicated ISGs, with expression levels similar to or greater than that of the myeloid cells in the tumor samples.

**Figure S8.**
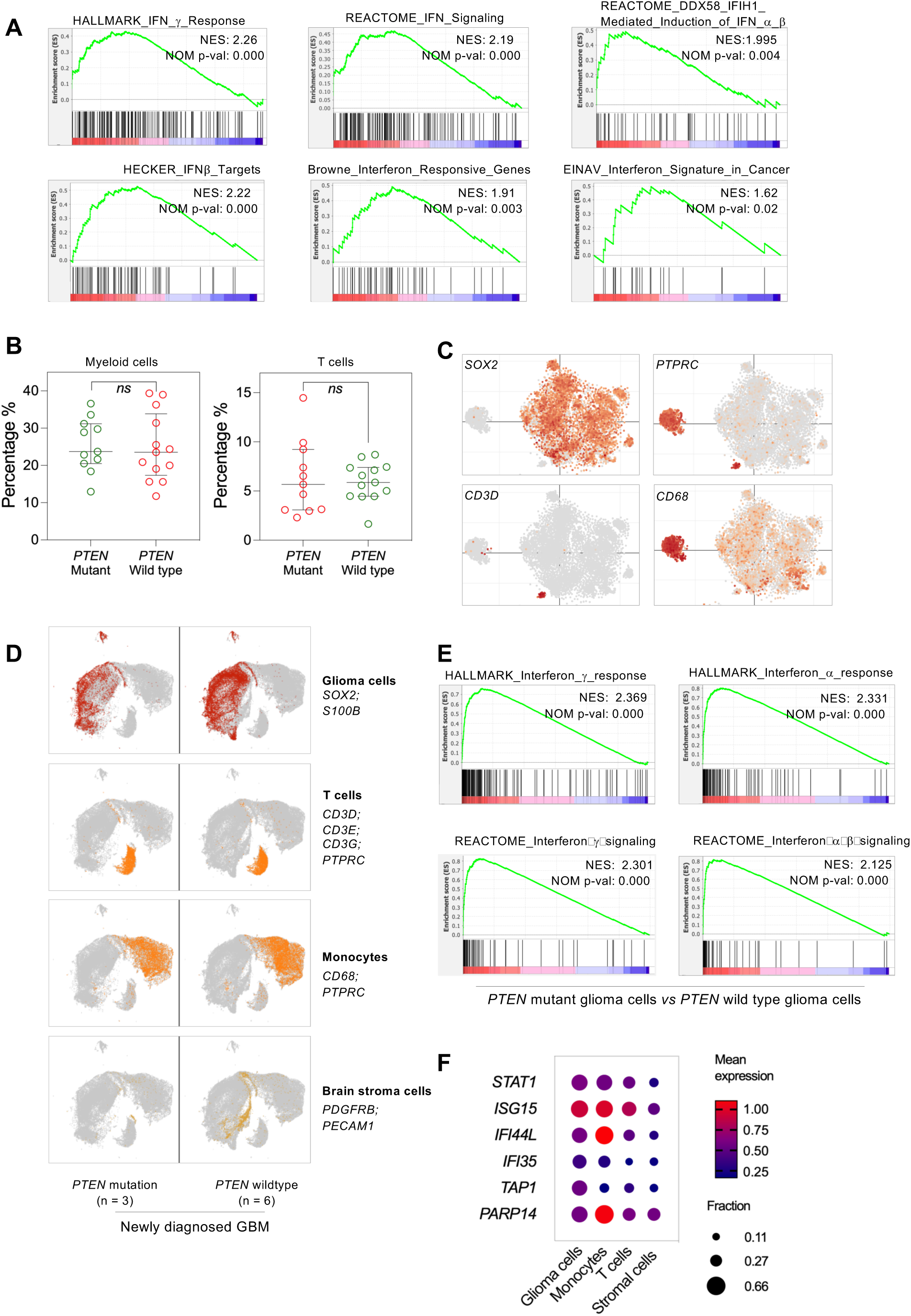
PTEN-mutated glioblastoma exhibits a robust activation of interferon response. (A) GSEA plots of the indicated interferon signatures for genes differently expressed in PTEN-mutated glioblastoma compared to PTEN wild type samples. Bulk RNA-seq data (SRA PRJNA482620) were downloaded and analyzed using kallisto and DESeq2. (B) Composition of myeloid and T cells in the glioblastoma cohort (Zhao et al., 2019). The analysis was based on bulk RNA-seq of tumor samples and performed via immune deconvolution using quanTIseq (Finotello et al., 2019). (C) t-SNE plots depicting the expression of the indicated genes in human glioblastoma (Neftel et al., 2019). Note that, compared to cancer cells (marked by SOX2 expression), macrophages (marked by the expression of CD68 and PTPRC) and T cells (marked by CD3D expression) comprise a smaller portion of the tumor mass in these samples. The plots were generated through queries and analysis of data available at the Broad Institute’s Single-Cell Portal (https://singlecell.broadinstitute.org/single_cell). (D) UMAP plots depicting different cell types present in PTEN-mutated or wild type glioblastoma samples. The indicated markers were used to identify these cell types. (E) GSEA plots of the most enriched interferon signatures for genes differentially expressed in tumor cells from PTEN-mutated glioblastoma compared to PTEN wild type. (F) For several ISGs, the mean expression and fraction of expressing cells in each of the four cell types in PTEN-mutated GBM samples were plotted. Note that tumor cells express these ISGs as robustly as immune cells within the tumor microenvironment.

**Figure S9.**
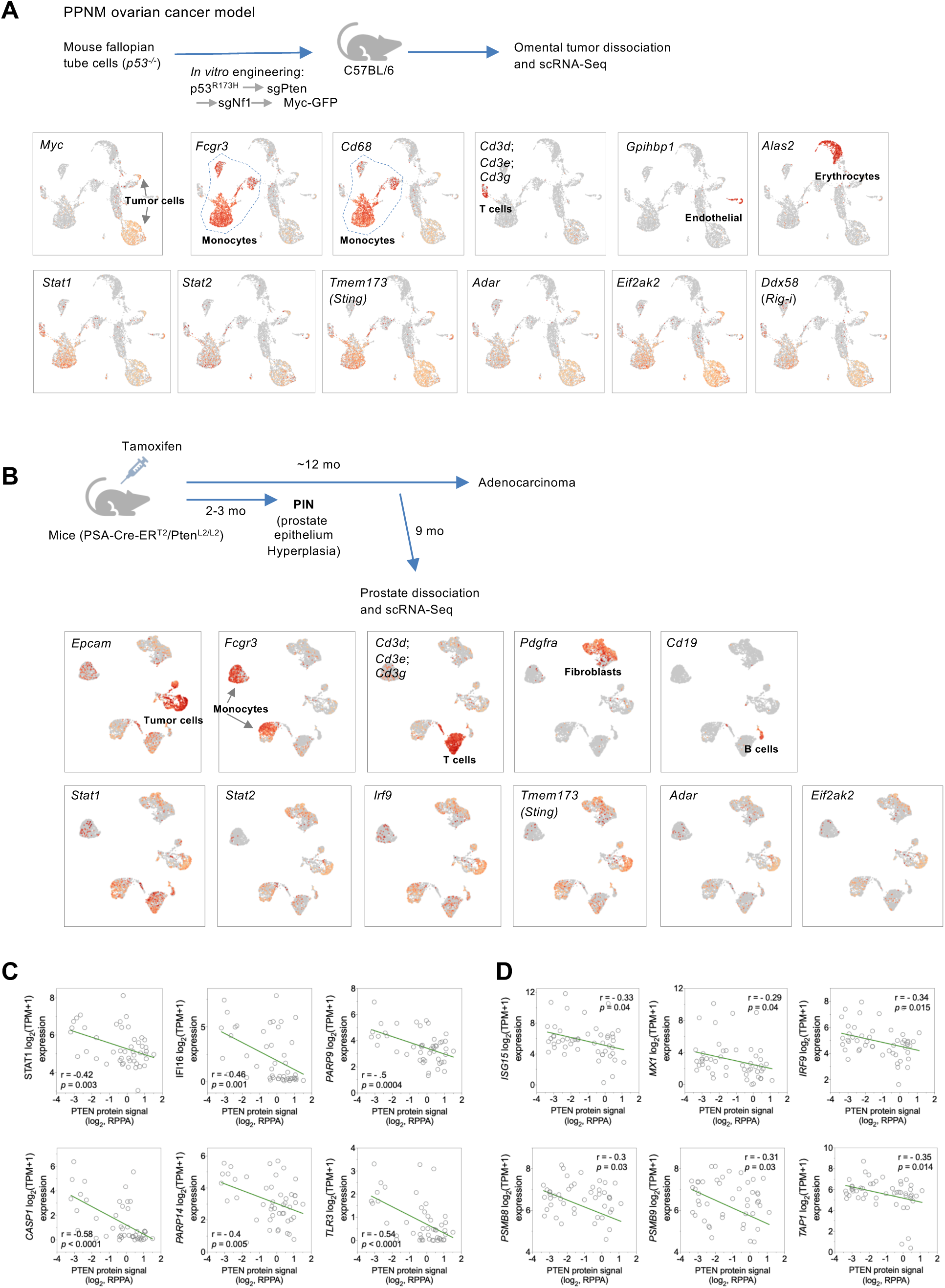
Mouse tumors derived from *Pten* loss exhibit tumor cell-intrinsic expression of interferon-stimulated genes. (A) (Top) Schematic diagram illustrating the derivation of the PPNM (loss of *T**<u>p</u>** 53*, ***<u>P</u>****ten*, ***<u>N</u>****f1*, and ***M****yc* overexpression) ovarian cancer model and tumor harvesting for scRNA-seq. The diagram was prepared using information from the original study (lyer et al., 2021). (Bottom) scRNA-seq data were downloaded from GEO GSE158474 and analyzed using the CellRanger pipeline to generate UMAP plots showing the expression of the indicated genes in different cell types. Note that ovarian tumor cells (marked by *Myc* expression) have robust expression of interferon-stimulated genes (ISGs), often similar to that of monocytes or T cells within the tumor microenvironment. (B) (Top) Schematic diagram illustrating Pten loss-driven prostate tumorigenesis (Abu el Maaty et al., 2021; Ratnacaram et al., 2008). (Bottom) UMAP plots of single-cell expression for the indicated genes. Note that prostate cancer cells (marked by *Epcam* expression) express ISGs at levels similar to those of immune cells within the tumor microenvironment. scRNA-seq data were downloaded from GEO GSE164858 and analyzed using the CellRanger pipeline. (C-D) Expression data for the indicated ISGs and PTEN protein abundance (determined by reverse-phase protein array) were downloaded from the Dependency Map (DepMap) portal (https://depmap.org/portal/) and analyzed in GraphPad Prism. Plots are shown for breast cancer cell lines (**C**, n = 47) and central nervous system (CNS) cancer cell lines (**D**, n = 50). Statistically significant correlation was not observed in cell lines of some other cancer types, likely due to the limited number of cell lines available for certain cancer types in the dataset and/or the distribution of PTEN protein abundance (e.g., lung cancer cell lines total 168 in the dataset, but their PTEN protein abundances are not evenly distributed as those of breast cancer cell lines).

**Figure S10.**
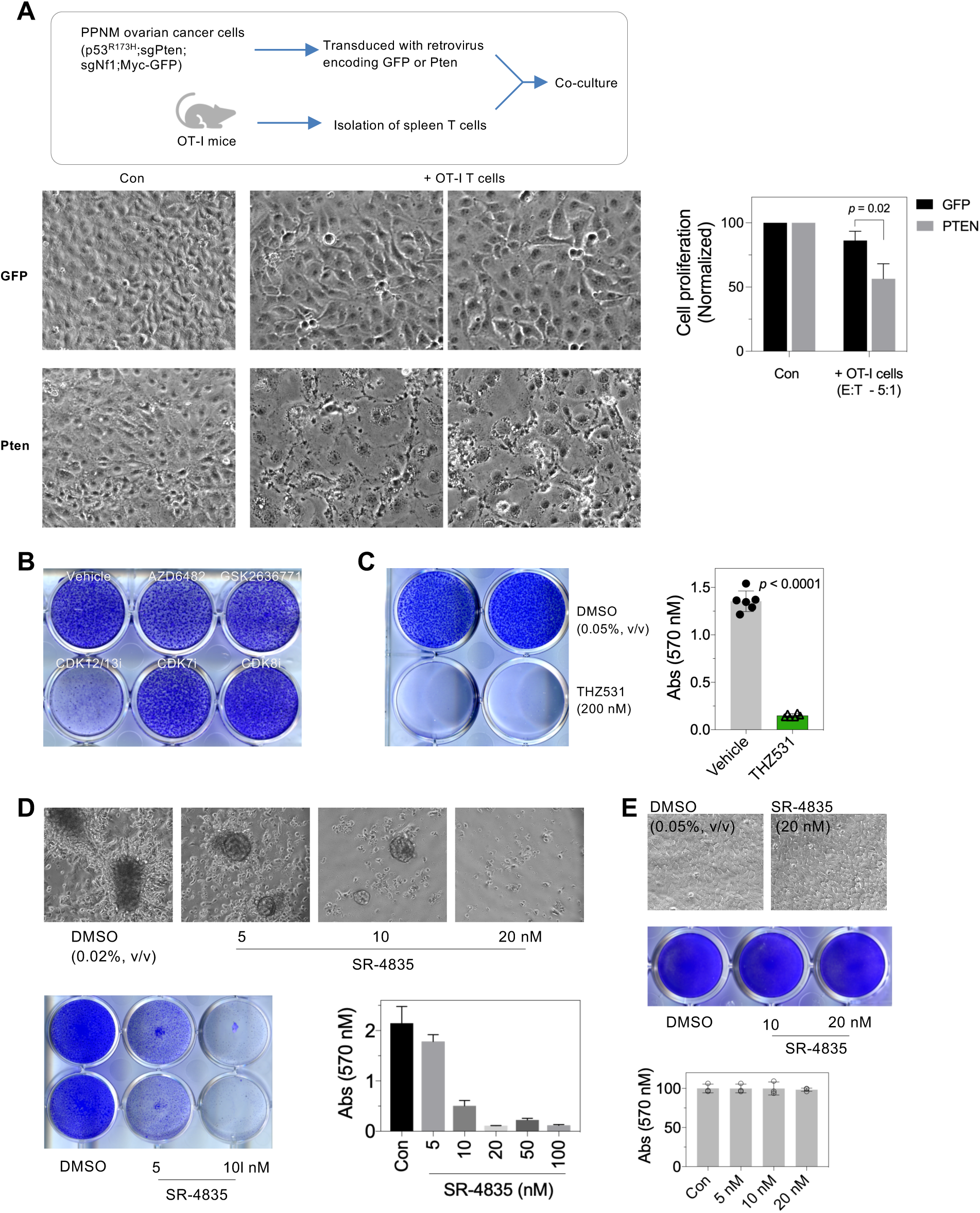
Assessing the response of PTEN-deficient cancer cells to cytotoxic immune cells and inhibition of transcriptional CDKs. (A) (Top) Schematic illustrating tumor cell – T cell co-culturing. (Bottom) PPNM cells expressing GFP control or Pten were pulsed with Ova (1 μg/ml final concentration) for 45 min, and seeded in 12-well plates (10,000 cells per well) with or without OT-I T cells (effector-to-target ratio: 5:1). One week after incubation, T cells were washed away, and cells were imaged using a 10x objective lens, followed by quantification of crystal violet staining. (B) MCF10A_sgPTEN cells were induced for transformation after one month of starvation of serum, growth factors, and insulin. The resulting colonies were treated with vehicle, PIK3CB inhibitors (1 μM KIN193/AZD6482 or 1 μM GS2636771), CDK12/13 inhibitor THZ531 (200 nM), CDK7 inhibitor YKL-5-124 (100 nM), or CDK8 inhibitor (200 nM CCT251545). The cultures were fixed after eleven days and stained with crystal violet. (C) MCF10A_sgPTEN cells were starved of serum, growth factors, and insulin. One week after the initiation of starvation, cells were treated with vehicle (0.05% DMSO, v/v) or THZ531 (200 nM). The cultures were fixed after 4 weeks for staining and quantification. (D) Cells were treated as in (B) with vehicle or CDK12/13 inhibitor SR-4835 at the indicated concentrations. Cells were imaged using a 10x objective lens (top), stained with crystal violet (bottom left), followed by quantification (bottom right). (E) MCF10A_sgPTEN cells were seeded at the density of 10,000 cells *per* well in 12-well plates in the presence of vehicle or SR-4835. After six days of treatment, cells were harvested for bright-field imaging (top), crystal violet staining (middle), and quantification (bottom).

## Materials and Methods

### Chemicals and Cytokines

PIK3CB inhibitor AZD6482/KIN193, PIK3CA inhibitor BYL719, and pan-PI3K inhibitor GDC-0941 were available from the central lab stocks that had been purchased as bulk orders from MedChemExpress. PIK3CB inhibitor GSK2636771 was purchased from MedChemExpress (5 mg size, #HY-15245). Recombinant Human IFN-α2 (carrier-free) and recombinant Human IFN-γ (carrier-free) were purchased from Biolegend (#592702 for IFN-α, #570206 for IFN-γ). Recombinant Human IFN-β was purchased from PeproTech (#300-02bc-5UG).

### Antibodies

Antibodies purchased and used for fluorescence immunoblotting are listed in the table below:

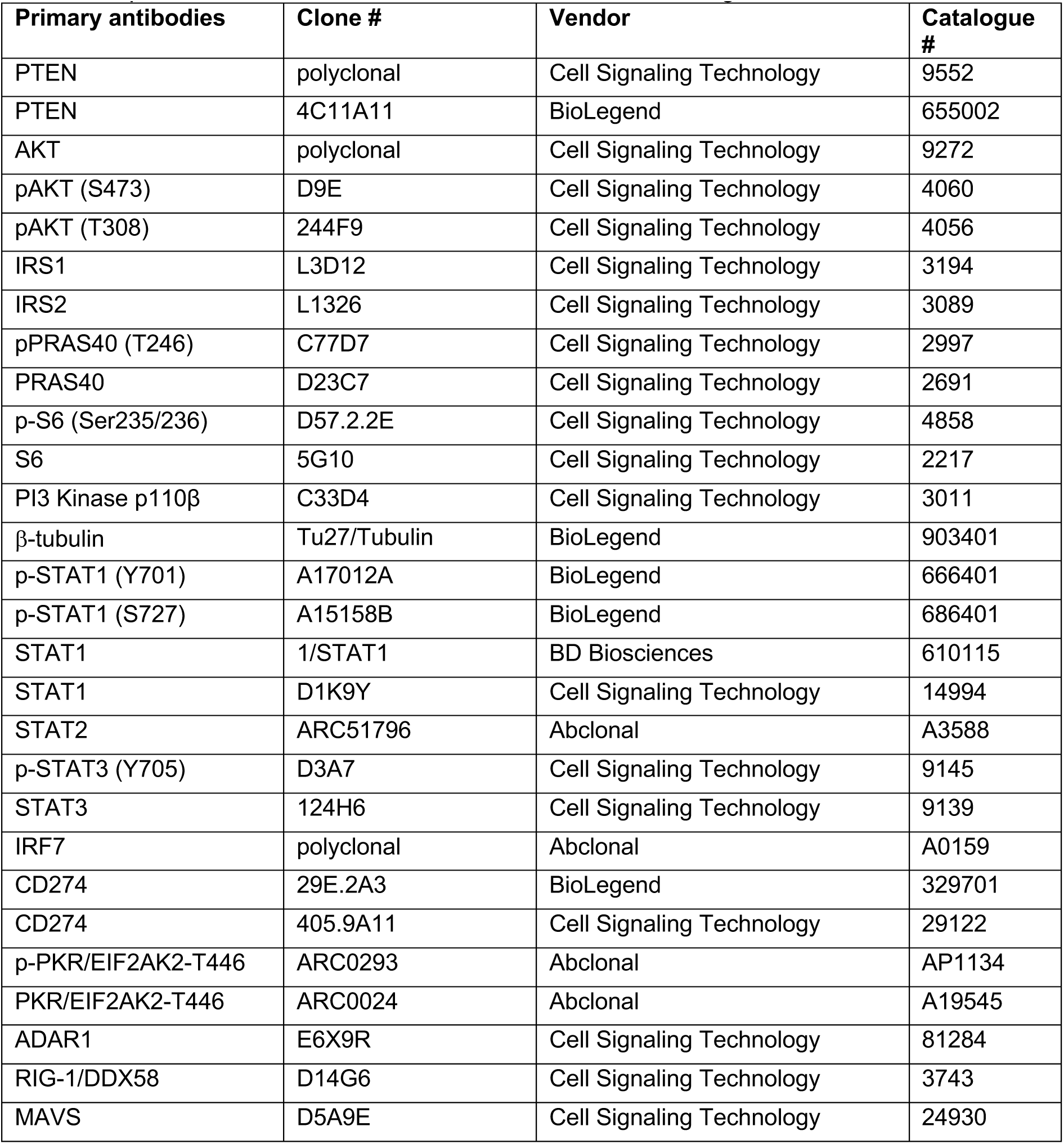
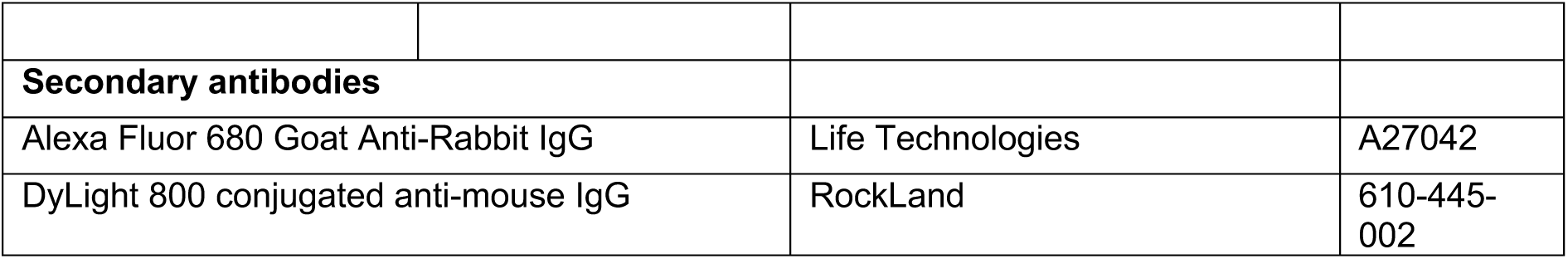

### Fluorescent Immunoblotting

Unless otherwise noted, cells were lysed with SDS sample buffer (50 mM Tris–HCl, pH 6.8; 2% (w/v) sodium dodecyl sulfate (SDS), 0.04% (w/v) bromophenol blue, 10% (v/v) glycerol, and 5% (v/v) β-mercaptoethanol). The amount of SDS sample buffer is typically 200 μl for one well of a 12-well plate seeded with 0.2 million human cells the day prior. Following collection, cell lysates were boiled for protein denaturation as well as sheering of the genomic DNA. Cell lysates were then loaded into SDS–polyacrylamide gel electrophoresis (SDS-PAGE) gels that were freshly cast.

Upon the completion of electrophoresis and transfer, the nitrocellulose membranes (Thermo Scientific, #88018) were blocked with Blotting Grade Blocker Non-fat Dry Milk (5%, w/v; Bio-Rad, #1706404XTU) and incubated with primary antibodies overnight in a cold room. The membrane was then washed and incubated with fluorescent-dye conjugated secondary antibodies for 1 h at room temperature. Following extensive washing, the membranes were scanned using an Odyssey CLx Near-Infrared Fluorescence Imaging System according to the manufacturer’s instructions. The images were exported and processed in ImageJ (NIH) for cropping and rotation before being imported into a PowerPoint file for assembly of blot images.

The protein marker used for immunoblotting is the PageRuler Plus Prestained Protein Ladder (Thermo Scientific, #26619). The markers emit near-infrared fluorescence signal in the 700 nm channel of the Odyssey CLx Infrared Imaging System and therefore provide critical information on protein size and the validity of the blotting signal. For use, the makers were diluted 20-fold in SDS sample buffer, and 20 μl of the diluted makers were loaded into one lane of the SDS-PAGE gels.

### Molecular Cloning

The lenti-viral constructs, pCDH-empty vector (pCDH-CMV-MCS-EF1α-Puro; System Biosciences, #CD510B-1), pCDH-PTEN, pCDH-CD274, were made following standard protocols of molecular cloning. Specifically, the templates used for amplifying the inserts were purchased from TransOmics (# BC074984, TCH1003 -MGC premier cDNA clone for CD274), InvivoGen (#puno1-mpten for PTEN). Primers for amplifying the inserts were synthesized at Eton Bioscience (Boston, MA), and PCR was performed using the KOD DNA polymerase (EMD Millipore). Purified insert DNA, as well as vector DNA, were digested with restriction enzymes (New England Biolabs, Ipswich, MA), followed by purification and ligation using T4 ligase (NEB, Ipswich, MA). Ligation product were then used to transform Stbl3 competent E.coli following standard procedures. After overnight incubation in the warm room, individual colonies were picked for PCR for identification of insertion, with positive colonies inoculated for culture in 25-50 ml LB medium supplemented with ampicillin. Plasmid DNA were extracted using QIAGEN Plasmid Plus Kits (Qiagen, #12943), and subjected to sequencing (Eton Bioscience, Boston, MA) for further validation. Plasmid DNA with fully sequenced insert were used for viral packaging and infection.

Lenti-viral CRIPSR vector (#49535, Addgene, Cambridge, MA) were used to introduce oligonucleotides for targeting genes of interest. Forward and reverse oligonucleotides were synthesized at Eton Bioscience and subjected to annealing, followed by ligation with a BsmBI-digested lentiCRISPR backbone. The ligation reaction was used to transform Stbl3 competent E. coli. Positive colonies that grew following overnight incubation in a warm room – identified by colony PCR using a U6 promoter paired with the reverse oligo of the guide insert – were subjected to further culture for plasmid DNA extraction (QIAGEN Plasmid Plus Kits, #12943). Plasmids were verified by sequencing (Eton Bioscience). Guide sequences were as follows: sgAAVS1 (also named sgCon in the text): 5’-GTCCCCTCCACCCCACAGTG-3’ sgPTEN_1 (also named sgPTEN in the text): 5’-TCATCTGGATTATAGACCAG-3’ sgPTEN_2: 5’-GGTTTGATAAGTTCTAGCTG-3’ sgSTING: 5’-AGAGCACACTCTCCGGTACC-3’ sgMAVS: 5’-CCTCTCCTGGAACTTCCGGT-3’

### Virus Packaging and Transduction

To package lenti-virus, HEK293T cells were seeded in T-25 tissue culture flasks (2.5 - 3 million cells *per* flask). Transfection was performed on the next day. Typically, 4 µg DNA including 2 µg vector DNA, 1.5 µg pCMVdR8.91, and 0.5 µg pMD2-VSVG were used and diluted in PBS. Polyethylenimine (PEI; homemade with powder purchased from Polysciences, # 23966-2) was also diluted in PBS before being mixed with diluted DNA for incubation (room temperature, 15 min).

Twenty-four hours after adding the lipid/DNA mixture to HEK293T cells, the cells were refreshed with new medium. Forty-eight and then seventy-two hours post the initial transfection, viral supernatant were collected and filtered through 0.45-µm filters and immediately used for transducing target cells supplemented with polybrene at a final dose of 8 µg/ml (Millipore, # TR-1003-G). Forty-eight hours after the initial infection, transduced cells were subjected to 2 days of puromycin selection (1-2 µg/ml, diluted from 10 mg/ml puromycin that was purchased from InvivoGen, #ant-pr-5). The cells were then subjected to a series of assays to examine the efficiency of gene editing as well as the potential effects on cell growth and their response to drug treatment.

### Cell culture

MCF-10A cells were from the stock of Roberts lab and originally purchased from the American Type Culture Collection (ATCC). Both MCF-10A_sgPTEN cells in regular culture and MCF-10_sgPTEN colonies were analyzed by STR (short tandem repeat) profiling (performed at DFCI Molecular Diagnostics Laboratory, using the GenePrint 10 System, Promega # B9510). All 10 STR marker genotypes had 100% match with published STR profiling (https://www.ncbi.nlm.nih.gov/biosample/?term=SAMN03471375)

Human breast cancer cell lines were also purchased from ATCC, and were cultured using RPMI1640 (Invitrogen, #11875) supplemented with 10% (v/v) fetal bovine serum (Gibco, #10438-026) and Penicillin-Streptomycin (Gibco, #15140122; a final concentration of 100 units *per* ml). Cells were incubated at 37°C with 5% CO_2_ and passaged twice a week using Trypsin-EDTA (0.05%) (Invitrogen, #25300). Cells were tested for mycoplasma contamination using the MycoAlert Mycoplasma Detection Kit (Lonza, # LT07-318), according to the manufacturer’s instructions.

### Cell Growth Assays

After trypsinization, cells were resuspended in medium for counting cell density using the Countess Automated Cell Counter (Life Technologies). Cells were seeded in 12-well plates at the density of 10,000 cells per well. Following one-week culture, cells were fixed and stained with crystal violet for visualization of cell growth. For quantification, the staining was extracted with 10% acetic acid, and absorbance measured at the 570 nm wavelength (750 nm used as the reference).

### Generation of MCF-10A_sgPTEN Colonies

MCF-10A cells with stable transduction of sgAAVS1 as control or sgPTEN_1 were seeded in 12-well plates at the density of 0.2 million cells per well. Following 3-4 days of culture, both cells reached visual confluency, and they were rinsed with DMEM/F-12 medium without any supplements or antibiotics and added with DMEM/F-12 medium for culture. Every 3-4 days, cells were refreshed with medium. After 4 weeks or longer, colonies became visually apparent in wells seeded with sgPTEN cells, while the majority of sgCon cells were dead and had been lost during medium refreshing.

### Correlation of drug sensitivity and gene dependency with PTEN status

Drug sensitivity data of 266 compounds from the GDSC project, genome-wide dependency scores from CRISPR (BROAD and Sanger) and RNAi (DRIVE and BROAD) screens, and Omics data were downloaded from DepMap. Cell lines were matched between the different datasets. Pearson correlations and their p-values were computed individually between the AUC scores of the entire drug panel and PTEN mRNA expression (n=533), copy number (n=755), mutation status (n=755) or protein expression (n=485). In the case of PTEN mutation status, the variable was binarized and a point biserial correlation was computed. Correlations and p-values were also determined between gene dependency scores and PTEN protein and, conversely, between PIK3CB dependency and the 214 probes in the RPPA dataset. All computations were performed using base packages in R. Correlation and volcano plots were generated in GraphPad Prism.

### RNA Extraction and Sequencing

Total RNA was prepared from tissue culture using the Monarch Total RNA Miniprep Kit (New England Biolabs, #T2010). On-column DNase digestion was included to remove any residual DNA. RNA integrity was assessed by an Agilent Bioanalyzer 2100 at the Novogene facility. The cDNA library was constructed using NEBNext® Ultra™ II RNA Library Prep with Sample Purification Beads (New England Biolabs, #E7775). Briefly, 400 ng RNA from each sample was used for poly(A)-selection, and then reverse transcription to generate cDNA. The resulting cDNA was subjected to adapter ligation and PCR amplification. Quality control of the library was performed using Labchip and qPCR. Sequencing was performed on a Novaseq 6000 to generate 150 bp paired-end reads (Novogene).

### RNA Sequencing Analysis

Raw reads containing adapters, base quality scores less than 5 and more than ten percent N’s were removed using Cutadapt v1.16. Processed reads were mapped to hg19 with STAR v2.7.3a 2-pass alignment using ENCODE parameters, first with a genome index of 100 bp overhangs and then with an index built using 75 bp overhangs and the splice junctions determined from the initial alignments. Unstranded gene hits were then counted for the alignments using HTSeq v0.9.1 with a minimum alignment quality of 10, and differential expression was determined in DESeq2. All genes having an average TMM-normalized CPM less than 1 were removed. Gene IDs were then annotated to their gene symbols and lowly expressed duplicates were also removed based on their average CPM. As independent validation, gene-level differential expression was determined from transcript abundances quantified in Kallisto v0.46 using the hg19 transcriptome. The differential expression analysis results of STAR and Kallisto were evaluated for correlation.

The RNA-seq data generated in this study has been deposited in the Gene Expression Omnibus (GEO) database (accession no. GSE229554).

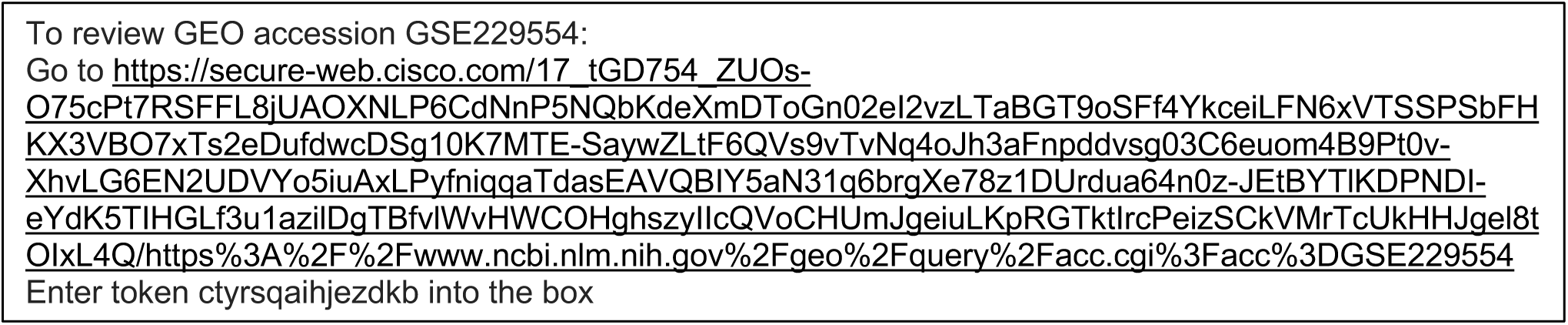

For GBM sample RNA-Seq (PRJNA482620), NCBI SRA-normalized reads were downloaded using fastq-dump and analyzed with Kallisto v0.46. Hg19 was used as the transcriptome assembly. Transcript abundances were quantified by applying default parameters and the strandedness information in the associated publication. Gene-level differential expression was then determined in DESeq2. For immune deconvolution of the bulk RNA-Seq, transcript-level counts from Kallisto were summarized into TPM-normalized gene-level abundances using tximport in R. The genome-wide abundances were then analyzed using quantTIseq to estimate the abundance of immune cells in the samples.

All volcano plots were plotted in PRISM using the log2 fold-change and adjusted p values determined with DESeq2. Significantly altered genes are those with |log2 FC| > 1 and q < 0.1. For gene set enrichment analysis, lists containing either all genes or only differentially expressed ones (q < 0.1) ranked by both the direction and statistical significance of their differential expression were inputted to GSEA v4.1.0 using default parameters and the Hallmark, Kegg, PID, Reactome and Biocarta pathway or gene set collections. Several other gene sets encoding interferon response genes and a recently identified gene set linked to cellular senescence (SenMayo), were also assessed for enrichment.

### TCGA Data Analysis

Singular Value Decomposition (SVD) was used to determine the most salient features from the RNA-Seq of the sgPTEN-transformed cells relative to control and pre-transformation sgPTEN cells. The top one percent of genes in each of the first two dimensions were merged to identify a gene signature associated with transformation. Then, raw counts of the TCGA breast cancer cohort (BRCA) were obtained from the UCSC Xena Data Browser and merged with that of the sgPTEN cells. The TCGA counts were generated in a workflow utilizing STAR 2-pass alignment with ENCODE parameters and counting with HTSeq. The raw counts were TMM-normalized and then log-transformed using edgeR v3.13. Genes in the derived signature were selected to generate a t-SNE plot using Rtsne v0.15 with a perplexity of 40.

### Single Cell RNA Seq Analysis

For GEM tumor sample data (GSE158886, GSE164858), NCBI SRA-normalized reads were downloaded from the European Nucleotide Archive and analyzed using Cell Ranger v5.0.1. A reference package was generated for mm10 using mkref, and fastq files were processed using the count pipeline. U-map projections were visualized in the loupe browser. The different cell types were determined based on commonly used markers and differentially expressed genes between cell clusters.

For GBM sample data (GSE182109), aligned BAM reads were downloaded from the European Nucleotide Archive. The downloaded bam files were converted to fastq using the bamtofastq tool. A reference package was generated for hg38 and the fastq files were processed using the count pipeline. The filtered featured matrices were further processed in R using the Seurat package. For each sample, cells with more than 500 features, less than 20% mitochondrial transcripts and less than 50% ribosomal transcripts and genes expressed in more than 5 cells were identified. Retained cell barcodes from each sample were pooled together, while retained genes were intersected between samples. The lists were stored in separate text files.

Samples were then aggregated using Cell Ranger, removing batch effects for the u-map and annotating for PTEN mutation status. Aggregated data were re-analyzed using the stored barcodes and genes. U-map projections were visualized in the loupe browser and all expression data were log-normalized. SOX2 and S100B were determined to be the most robust markers distinguishing glioma cells from immune cells and brain stroma. Therefore, epithelial cells were determined based on the following criteria: SOX2 > 0, S100B > 0, CD68 = 0, PTPRC = 0, PECAM1 = 0, PDGFRB = 0 and PDGFRA = 0. Differential expression was determined for epithelial cells in PTEN mutated vs wild-type samples. GSEA was performed on differentially expressed genes (q < 0.1) using the previously mentioned gene set collections. To quantify the expression of individual genes in a cell type, the fraction of cells expressing the gene and the gene’s average, log-normalized UMI counts were determined. Dot plots were generated in PRISM.

For the scRNA-Seq data generated in this study of the sgPTEN transformed cells, fastq files were processed using the Cell Ranger count pipeline and hg38 reference package. The filtered feature matrix was loaded into Seurat, and cells with more than 300 gene features, more than 500 UMI counts in total, less than 30% mitochondrial transcripts and more than 5% ribosomal transcripts were retained. The filtered data were processed using SCTransform to remove genes that were expressed in fewer than 5 cells. The unscaled, log-normalized counts were used to determine the Spearman rank correlation between STAT1 and other ISGs in cells expressing the two genes using base packages in R. Scatter plots were generated in PRISM.

